# The protein-protein interaction landscape of Fanconi anemia protein A (FANCA) reveals its contributions to mRNA processes

**DOI:** 10.1101/2025.09.11.675346

**Authors:** Virgile Gibert, Frédérique Maczkowiak-Chartois, Benedetta Mancini, Emilie-Fleur Gautier, Johanna Bruce, Morgane Le Gall, Serge Urbach, Angelos Constantinou, Anne Helbling-Leclerc, Jihane Basbous, Filippo Rosselli

**Affiliations:** Université Paris-Saclay, CNRS-UMR9019 Genome Integrity and Cancers, Gustave Roussy, Villejuif, France; ENS Paris-Saclay, Gif-sur-Yvette, France; Proteom’IC facility, Université Paris-Cité, CNRS, INSERM, Institut Cochin, Paris, France; Institut de Génomique Fonctionnelle, Université de Montpellier, CNRS, INSERM, Montpellier, France; Montpellier RIO Imaging, Montpellier, France; Institut de génétique Humaine, Université de Montpellier, CNRS, Montpellier, France

## Abstract

Fanconi Anemia (FA) is a rare inherited recessive disorder characterized by bone marrow failure, congenital abnormalities, and a predisposition to myeloid leukemia and some solid cancers. FA is caused by mutations in one of the 22 so-called FANC genes, with two thirds of FA patients being FANCA mutants. FANC proteins cooperate within the FANC/BRCA pathway, a DNA repair system essential for resolving DNA interstrand crosslinks (ICLs) in S-phase. The canonical understanding of FA pathogenesis involves genome instability and impaired cellular proliferation due to DNA damage accumulation and stress response overactivation. However, recent data shows FANCA has a function in ribosome biogenesis and translation, which prompts a reassessment of FANCA’s role(s) in cellular physiology and FA pathogenesis. We reasoned that unveiling the multiple interactions of the FANCA protein would shed light on these potentially new FANCA functions. By combining immunoprecipitation and BioID approaches, in three different cell lines and with different in silico analysis methods, we constructed lists of FANCA interactome proteins of different levels of selectivity and assembled them into a novel multilayered FANCA protein-protein interaction (PPI) landscape. Strikingly, we find that FANCA associates with proteins involved in processes distinct from DNA repair, such as translation, ribosome biogenesis, mRNA splicing, nucleocytoplasmic transport, chromatin remodeling, and the TCA cycle. We then analyzed the protein content of polysome profiling fractions and found that FANCA loss may affect ribosomal protein stoichiometry within ribosomal subunits. Overall, our data expands FANCA’s PPI network far beyond the previously reported interactions. Importantly, we confirm the biological significance of low-selectivity interactions and propose to re-evaluate their importance in proteomic studies. The study of previously unsuspected FANCA functions, such as translation, could provide new therapeutic targets in FA and therefore improve patient care.

## Introduction

Loss-of-function mutations in more than 20 genes (*FANCA* to *FANCW*) have been identified as the genetic basis of Fanconi anemia (FA), a rare inherited bone marrow failure syndrome characterized by progressive quantitative and/or qualitative defects in hematopoietic stem and progenitor cells (HSCs/HSPCs), leading to pancytopenia, anemia, and thrombocytopenia. Additionally, FA patients often present with various physical and functional anomalies, including skeletal malformations (particularly affecting the head, forearm, and thumb), skin pigmentation changes (e.g., café-au-lait spots), endocrine disorders (such as diabetes), sensory deficits (e.g., deafness), infertility, immune dysfunction, and cognitive impairment. Despite the clinical complexity and heterogeneity of the syndrome, approximately 20% of FA patients are asymptomatic at birth. FA patients are also at risk of developing myelodysplastic syndrome (MDS), acute myeloid leukemia (AML), and solid tumors (1–5).

The FANC proteins form several biochemical modules that act sequentially along the FANC/BRCA-homologous recombination (HR) and replication rescue DNA repair pathway (FANC/BRCA pathway) (6), an integral part of the broader DNA damage response (DDR) network which contributes to genome integrity maintenance, thereby relieving cells from genotoxic and replicative stress. Inside the pathway, the FANCcore complex links the sensing of replication stress (through FANCM) to the HR-mediated rescue of stalled/delayed replication forks *via* FANCD2 and FANCI monoubiquitination (see Figure 2A). Disruption of the FANC/BRCA pathway results in cellular and chromosomal hypersensitivity to DNA interstrand crosslinks (ICLs) and replication stress (7–10). Additionally, beyond its canonical role in DNA repair, the pathway contributes to various genome stability mechanisms, including protection of common fragile site (9, 11, 12), regulation of R-loop homeostasis (13, 14), telomere length maintenance (15, 16) and proper mitotic progression (17–19). Moreover, FA cells show several other cellular and molecular abnormalities, such as a pro-oxidative and pro-inflammatory cellular environment, and dysregulated cytokine and growth factor signaling involving Tumor Necrosis Factor-α (TNF-α), Transforming Growth Factor-β (TGF-β) and Interferons (IFNs) (20–24). FA cells also exhibit mitochondrial dysfunction (25, 26) and impaired proteostasis (4, 27, 28). Given the extensive heterogeneity of FA phenotypes across patients and cell types, it is plausible that defects in processes distinct from DNA repair account for the full spectrum of the disease. Indeed, the FANC/BRCA pathway, or specific components of it, may have alternative molecular or biochemical functions beyond the DDR.

To explore this hypothesis, we aimed to define the protein-protein interaction (PPI) network of FANCA, the most frequently mutated gene among FA patients, responsible for approximately 60-70% of FA cases worldwide. Our analysis combined data from three approaches: mass spectrometry (MS)-based identification of proteins (a) co-immunoprecipitating with endogenous FANCA in two distinct cell lines; (b) biotinylated by a FANCA-BirA fusion protein expressed in a third cell line; and (c) analysis of previously published interaction data. This integrative strategy allowed us to construct a comprehensive FANCA PPI landscape encompassing both known and novel interactors.

Our results reveal that FANCA interacts with proteins involved in a wider range of processes than genome integrity maintenance. Specifically, FANCA appears to participate in pathways involved in multiple aspects of mRNA metabolism, including splicing, nucleocytoplasmic transport, and translation, suggesting a contribution to a broader spectrum of cellular functions than previously thought, and which may underlie the complex phenotypic presentation of FA.

## Results

### Generation of FANCA protein-protein interaction datasets

We aimed to obtain a comprehensive overview of the FANCA protein–protein interaction (PPI) network, encompassing physical and indirect, as well as vicinity and functional associations. To this end, endogenous FANCA was immunoprecipitated (IP) from whole-cell extracts of HSC93 lymphoblasts, an EBV-immortalized cell line derived from a healthy donor and widely used as a standard control in in vitro FA studies. Co-immunoprecipitated (co-IP) proteins were identified and quantified via label-free quantitative MS.

We conducted four independent IP experiments using HSC93 cell extracts (Figure 1A). To reduce nucleic acid–mediated artifacts, cell extracts were treated with benzonase, a nuclease that digests both RNA and DNA, and subsequently sonicated. IP was then performed using an anti-FANCA antibody or an isotypic IgG control. Across these experiments, MS analysis identified between 720 and 3,234 proteins per IP, resulting in a total of 3,420 unique proteins across all FANCA and control IPs (Supplemental Table (ST)1, Sheet 1). For each protein, a label-free quantification (LFQ) value was determined in each experiment.

**Figure 1.**
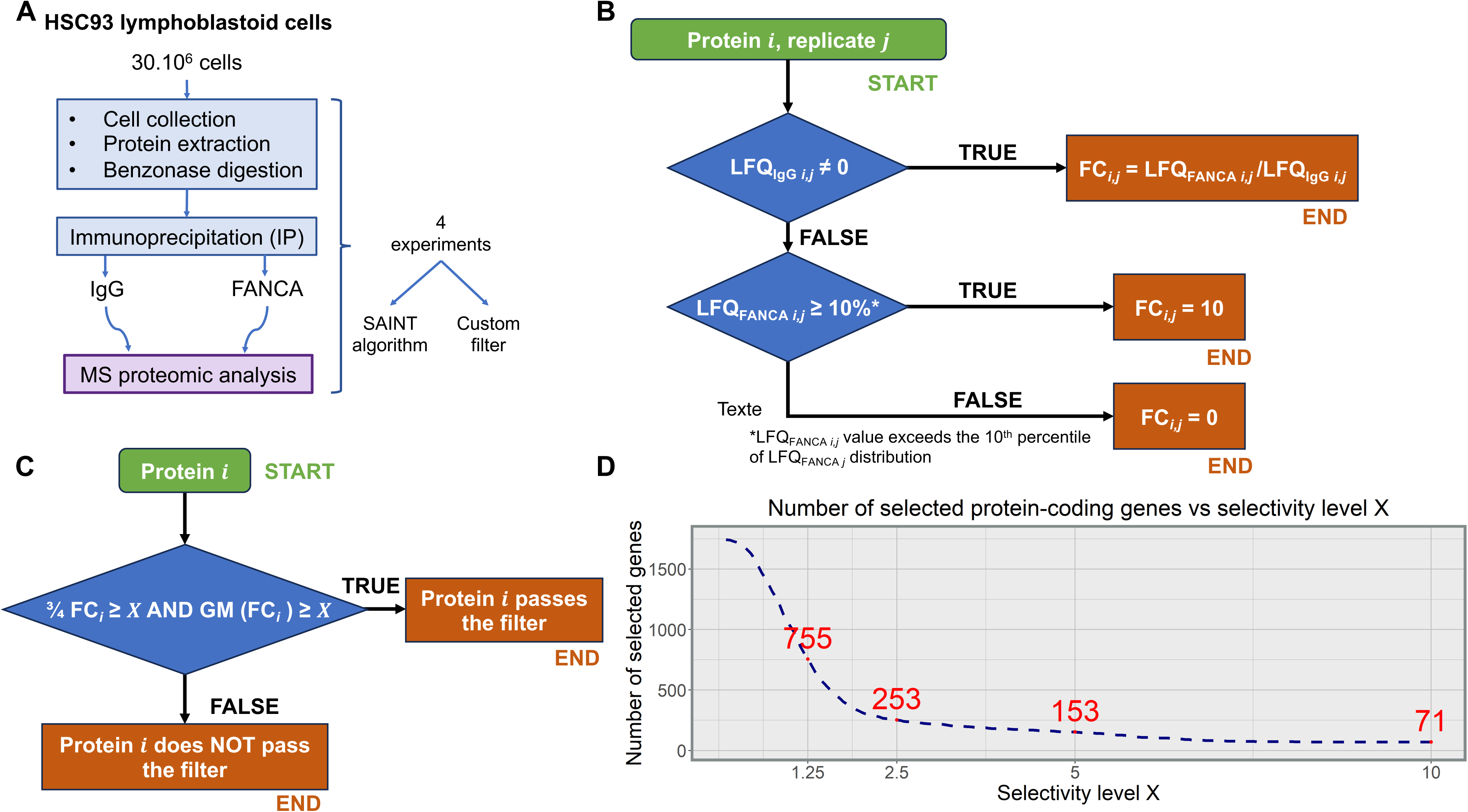
FANCA immunoprecipitation in HSC93 cells: pipeline of analysis and data filtering. **(A)** Schematic representation of our experimental and analytical procedure. **(B)** IP FANCA/ IP IgG ratio calculation algorithm. **(C)** In-house interactor filter used with HSC93 IP data. **(D)** Number of genes passing the filter as a function of selectivity level **X**=1.25, **X**=2.5, **X**=5, **X**=10.

**Figure 2.**
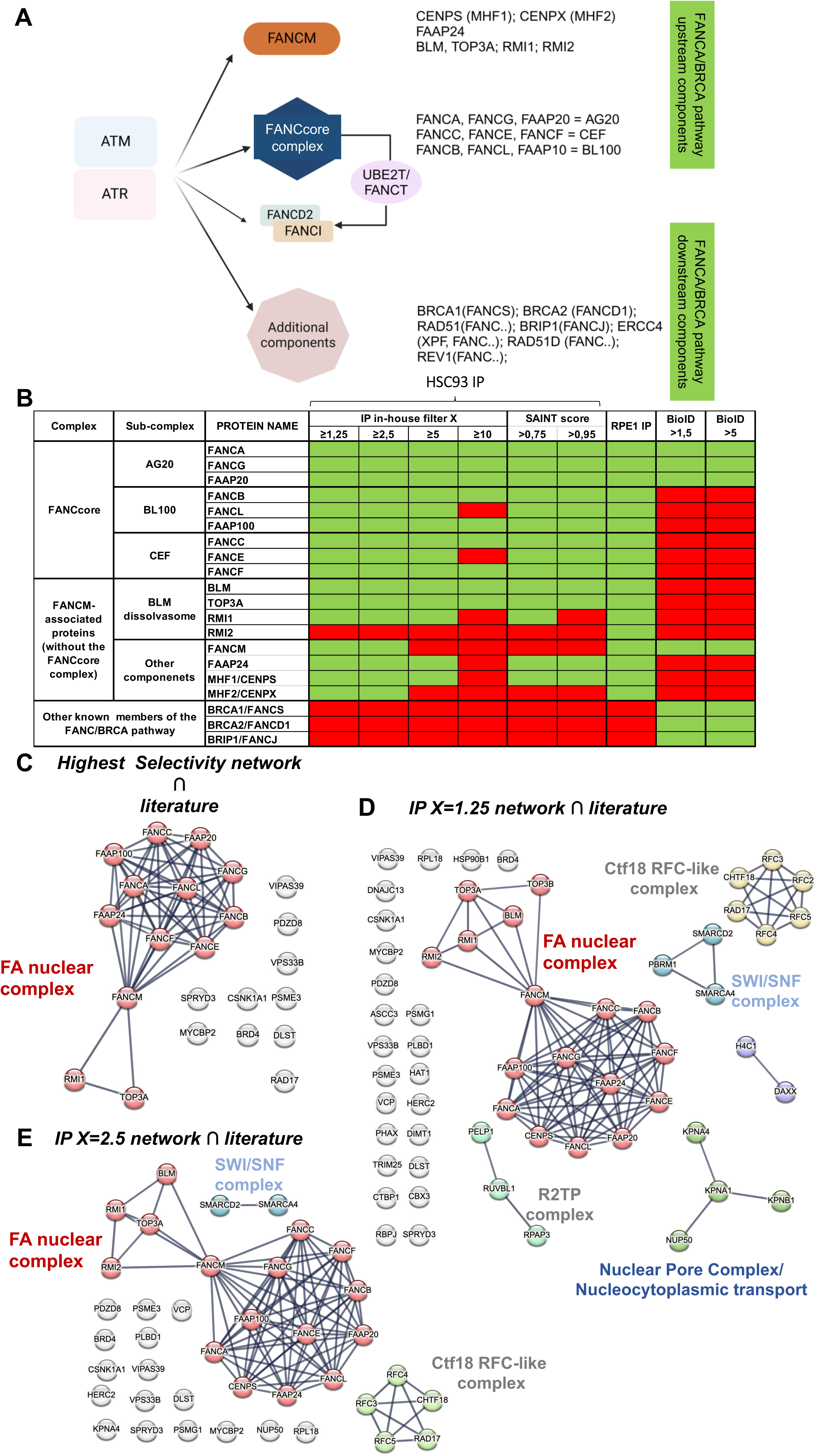
Common preys of FANCA between our analysis and previously published works. **(A)** Schematic representation of the FANC/BRCA pathway. As indicated, the FANC core complex consists of the assembly of three subcomplexes, AG20, CEF and BL100, each of three proteins. It acts as an E3 ubiquitin ligase that catalyzes, via FANCL, the transfer of a ubiquitin moiety from the E2 enzyme UBE2T/FANCT to its two substrates: FANCD2 and FANCI. The monoubiquitination of both FANCD2 and FANCI is required for their assembly into chromatin-associated nuclear foci to guide the recruitment of the FANCA/BRCA pathway downstream components leading to homologous recombination. **(B)** FANCA interactions within the FANCA/BRCA pathway proteins identified by MS analysis in co-IP or BioID extracts from HSC93, RPE1 or HEK cells as revealed by different analytical approaches. **(C to E)** Clustering of common FANCA preys between literature and our IP datasets. Clusters were proposed by STRING on the basis of the following setting: network type’physical subnetwork’; meaning of network edge’confidence’; minimum required interaction score’highest confidence’ (0.9); for C and D,’hide disconnected node in the network’; MCL clustering with an inflation parameter of 3 without edges between the clusters.

Two complementary in silico strategies were employed to prioritize potential FANCA interactors. The first used SAINT (Significance Analysis of INTeractome), a probabilistic scoring algorithm specifically designed for IP-MS data, which assesses the likelihood of true biological interactions (29). This analysis yielded two sets of FANCA-interacting proteins based on SAINT probability (SP) thresholds: 108 proteins with SP>0.75 and 58 proteins with SP>0.95 (Table ST1, Sheets 2 and 3; Supplemental Figure (S)1A).

We developed a second filtering method, hereafter referred to as the in-house filter. Each protein (i) is assigned a fold change (FC) ratio in each experiment (j), calculated as:

FC***_i_***_,j_=LFQ_FANCA,***i***,j_/LFQ_IgG,***i***,j_

For proteins undetected in the IgG control (LFQ_IgG,***i***,j_=0) FC***_i_***_,j_ is set to 10, provided that LFQ_FANCA,***i***,j_ exceeds the 10^th^ percentile of LFQ_FANCA,j_ values in that experiment. Otherwise FC***_i_***_,j_ is set to 0 (Figure 1B). To filter for potential interactors, a protein is retained at a given selectivity level **X** if at least three out of four FC values (from the four experiments) exceed **X**, and if the geometric mean of all four FC values is also greater than **X**. The geometric mean criterion reduces the influence of outliers and enhances selectivity (Figure 1C). The filtering is applied at four selectivity levels: **X**=1.25, 2.5, 5.0, and 10, yielding interactor lists containing 756, 253, 153, and 71 proteins, respectively (Figure 1D; Table ST2, Sheets 1–4).

Finally, we generated a high-confidence list of FANCA interactors by combining proteins identified at the highest selectivity level (**X**=10) with those scoring above SP>0.95 in the SAINT analysis. This approach yielded a total of 93 unique proteins (Table ST3, Sheet 1). In total, we generated seven candidate FANCA partner lists to be used for subsequent analysis (Table ST3, Sheet 1).

### Validation of the identified FANCA partners: a literature analysis

Before proceeding with the in silico analysis of the biological functions associated with the proteins we co-IP with FANCA, we assessed the reliability of our strategy by comparing our results with previously published datasets. To this end, we compiled a reference list of known FANCA interactors by aggregating the 159 proteins listed in the BioGRID database (version 4.4.240) (30) with the 134 proteins identified through co-IP and SAINT analysis (SP>0.90) in the HEK293 human embryonic kidney cell line (31) and some additional interactors reported in literature but not included in BioGRID dataset (27, 32, 33). This established a list of 316 potential FANCA preys (Table ST3, Sheet 2).

Among the 93 proteins in our high-confidence list of FANCA preys, 23 (24.7%) have already been reported as FANCA interactors (Table ST3, Sheet 3). We also assessed the overlap with known FANCA partners at lower selectivity levels of the in-house filter. At **X**=1.25, 59/757 proteins (7.8%) matched previously reported interactors; at **X**=2.5, 39/254 proteins (15.4%); and, at **X**=5, 30/154 proteins (19.5%) (Table ST3, Sheet 3). Expectedly, the overlap increases with selectivity.

Limiting our comparison to the 159 FANCA interactors listed in the BioGRID database, we identified 16 (10%) proteins among the 153 identified at selectivity level **X**=5 and 13 (14%) among the 93 high-confidence interactors (**X**=10 or SP>0.95). This is much more than in Lagundžin et al. (31), which retrieved only 7 (5%) of identified proteins registered in the BioGRID database in their dataset of 134 Co-IP proteins (SP>90) (Table ST3, Sheet 2).

Remarkably, among the range of proteins known to associate with FANCA found at **X**=2.5, we identify almost all of the upstream components of the FANC/BRCA pathway, i.e. the entire FANCcore complex and most key FANCM-associated partners (Figure 2A to E, Figure S1B), components of the SWI/SNF (SWITCH/Sucrose Nonfermentable) chromatin remodeling complex such as SMARCA4 (also known as BRG1) (32, 34), PBRM1 (BAF180) (35), and SMARCD2 (36), as well as several components of the RFC and RFC-like complexes, *i.e.* RFC2-5, RAD17 and CHTF18 (31, 36), the nuclear pore complex (NPC) protein NUP50 (31), the nuclear-cytoplasmic shuttles KPNA1, KPNA4 and KPNB1 (31, 32), three components of the R2TP complex, *i.e.* RUVBL1 (37), RPAP3 (31) and PELP1 (31), the chaperon HSP90B1 (38), and the ribosomal protein RPL18 (32) (Figure 2C to E; Table ST3, Sheet 3).

Overall, these analyses demonstrate the consistency of our approach and the relevance of our datasets for in silico functional analyses. It is also important to emphasize that our main objective is to characterize the entire set of proteins influenced by FANCA, and not just to capture the most stable or abundant physical interactions.

### Validation of the identified FANCA partners: alternative experimental approaches

To support and extend our findings, we conducted two additional experiments using alternative approaches. First, we repeated FANCA IP in extracts from human RPE1 cells, an hTERT-immortalized and p53-proficient epithelial cell line. Second, we used a proximity-dependent biotin labeling strategy (BioID) coupled with MS analysis in Flp-In HEK293 cells expressing a FANCA-BirA fusion protein. Unlike IP, which relies on biochemical affinity, the BioID method identifies proteins in close spatial proximity to FANCA (i.e. its “proxilome”).

From IPs in RPE1 cells using anti-FANCA and isotype control IgG antibodies, we identified 303 candidate FANCA-associated proteins. The proteins retained are those detected with at least three unique peptides in the FANCA IP and no peptide in the IgG IP, matching the detection pattern shown by FANCA interactor FAAP20 (Table ST3, Sheet 1; Table ST4, Sheet 1). Among the 93 proteins identified in the most selective HSC93-derived lists (**X**=10 and SP>0.95), 43 (46%) were also detected in the RPE1-derived IP dataset (Table ST3, Sheet 4). At **X**=1.25, **X**=2.5, and **X**=5, 148, 75, and 48 proteins, respectively, are also present in the RPE1 IP (Figure 3A, C; Table ST3, Sheet 4).

**Figure 3.**
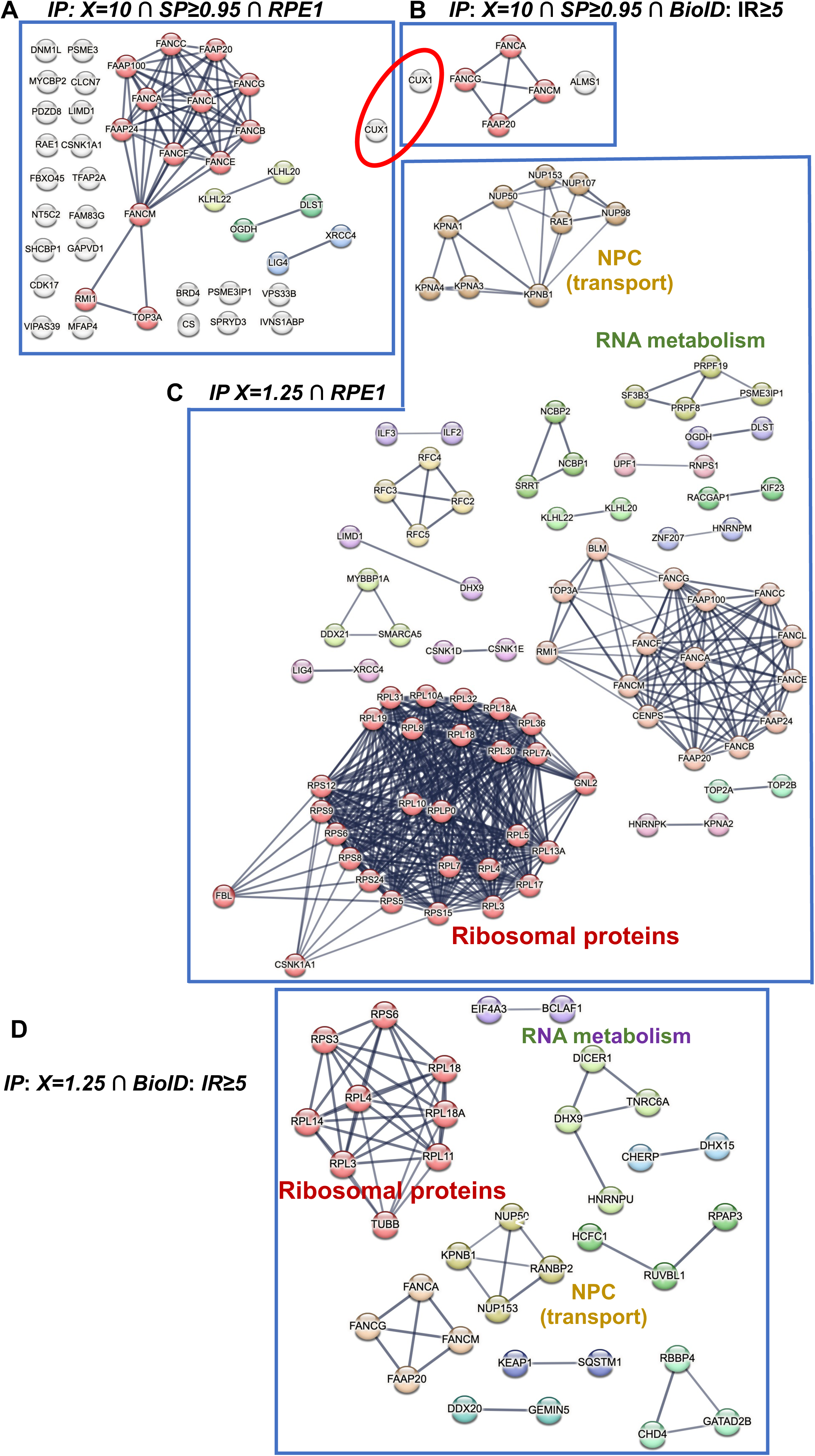
STRING based clustering of FANCA preys at different levels of selectivity as emerging from datasets obtained following IP or BioID analysis in different cells. **(A)** Proteins identified as FANCA partners in the **X**=10 and SP>0.95 lists and present in the RPE1 IP, or **(B)** in the BioID>5X analysis. **(C)** Clustering of FANCA co-immunoprecipitated proteins at **X**=1.25 and in RPE1, or **(D)** at **X** =1.25 and identified as biotinylated at selective level >1.5. Clusters were proposed by STRING on the basis of the following setting: network type’physical subnetwork’; meaning of network edge’confidence’; minimum required interaction score’highest confidence (0.9); for C and D,’hide disconnected node in the network’; MCL clustering with an inflation parameter of 3 without edges between the clusters.

The BioID-MS analysis yielded 344 biotinylated proteins (Table ST5, Sheet 1), from which we derived two subsets based on the intensity ratio (IR) between experimental and control samples: IR≥1.5 (217 proteins) and IR≥5 (133 proteins) (Table ST3, Sheet 1). As expected, given the methodological differences, the overlap between the IP and BioID datasets was more limited. Only 6 proteins were shared between our most selective HSC93 IP list (**X**=10 or SP>0.95) and the BioID datasets (Figure 3B). At IR≥1.5, 53, 17, and 8 proteins overlapped the HSC93 IP lists at **X**=1.25, **X**=2.5, and **X**=5, respectively (Figure 3D; Table ST3, Sheet 4). At IR≥5 level, these numbers decreased to 29, 8, and 6, respectively.

Despite originating from single experiments and therefore lacking statistical weight, these complementary approaches contribute to a more comprehensive view of the FANCA PPI network, capturing not only high-affinity interactors but also spatially and functionally relevant partners. Indeed, only four proteins, FANCG, FAAP20, FANCM, and CUX1, were consistently identified across all three platforms (HSC93 IP, RPE1 IP, and HEK293 BioID) at the highest selectivity levels (Figure 3A and B). CUX1 is a transcription factor involved in differentiation, cell cycle control, and DNA damage response, with potential tumor suppressor functions (39, 40). At lower selectivity thresholds, both IP and BioID datasets converge on several additional FANCA-associated proteins involved in translation, nucleocytoplasmic transport, and RNA metabolism (Figure 3C and D), suggesting FANCA may broadly influence gene expression (from transcription to translation) beyond its canonical role in genome maintenance.

### FANCA interactors link FANCA to genetic information processing and metabolic processes

To unravel the biological functions associated with FANCA’s PPI network, we conducted an in silico functional analysis of the protein lists generated by the in-house filter on the HSC93 IP data (Table ST2, Sheet 1-4, *i.e.* **X**=1.25 to **X**=10) using three functional term libraries: the Kyoto Encyclopedia of Genes and Genomes (KEGG, version 113.0) (41), the Gene Ontology Biological Process (GO_BP) and the Gene Ontology Cellular Components (GO_CC) (Release 2025-03-16) (42). The analysis was performed using the ClusterProfiler R package (43).

Starting at **X**=1.25, we identify 19 KEGG pathways, belonging to four categories, namely “Genetic Information Processing”, “Human Diseases”, “Cellular Processes”, and “Metabolism”, which are significantly enriched (Figure 4A; Table ST2, Sheet 5). Consistent with the role of FANCA in DNA repair and replication rescue, 6 of the 19 enriched terms belong to the subcategory “Replication and Repair”, namely “Fanconi Anemia Pathway”, “Mismatch Repair”, “Base Excision Repair”, “DNA Replication”, “Nucleotide Excision Repair”, and “Homologous Recombination” (Figure 4A; Table ST2, Sheet 5). Among the other “Genetic Information Processing” enriched terms, two are related to chromatin dynamics, namely “ATP-dependent chromatin remodeling” and “Polycomb repressive complex”, four are included in the subcategory “Translation”, i.e. “Ribosome” (the most enriched KEGG term at **X**=1.25, adjusted *p*-value=6.02×10^-60^), “Ribosome biogenesis in eukaryotes”, “Nucleocytoplasmic transport”, and “mRNA surveillance pathway”. Other mRNA-related terms are present: “Spliceosome”, “RNA polymerase”, and “RNA degradation” (Figure 4A; Table ST2, Sheet 5). At **X**=2.5, the enrichment in RNA-related terms no longer meets the *p*-value threshold; five new terms appear, all related to carbon metabolism (Figure 4A; Table ST2, Sheet 6). At **X**=5, 6 of the 11 retained terms are related to carbon metabolism. The others include “Fanconi anemia pathway”, “ATP-dependent chromatin remodeling”, “Homologous Recombination” and “Non-Homologous End Joining” (Figure 4A; Table ST2, Sheet 7). At the highest selectivity level studied, i.e., **X**=10, 5 of the 7 terms are related to carbon metabolism, the remaining two terms being “Non-Homologous End Joining” and “Fanconi anemia pathway”, which is the most significantly enriched at **X**=10 (*p*-value=09.3×10^-9^) (Figure 4A; Table ST2, sheet 8). Functional enrichment using GO_BP and GO_CC terms was consistent with the KEGG enrichment (Figure 4B-C; Table ST2, Sheets 9-16), with GO_CC term “Fanconi anemia nuclear complex” and GO_BP term “Interstrand crosslink repair” being significantly enriched at all selectivity levels, whereas the RNA processing-and translation-related processes/complexes are enriched at the lower **X** levels.

**Figure 4.**
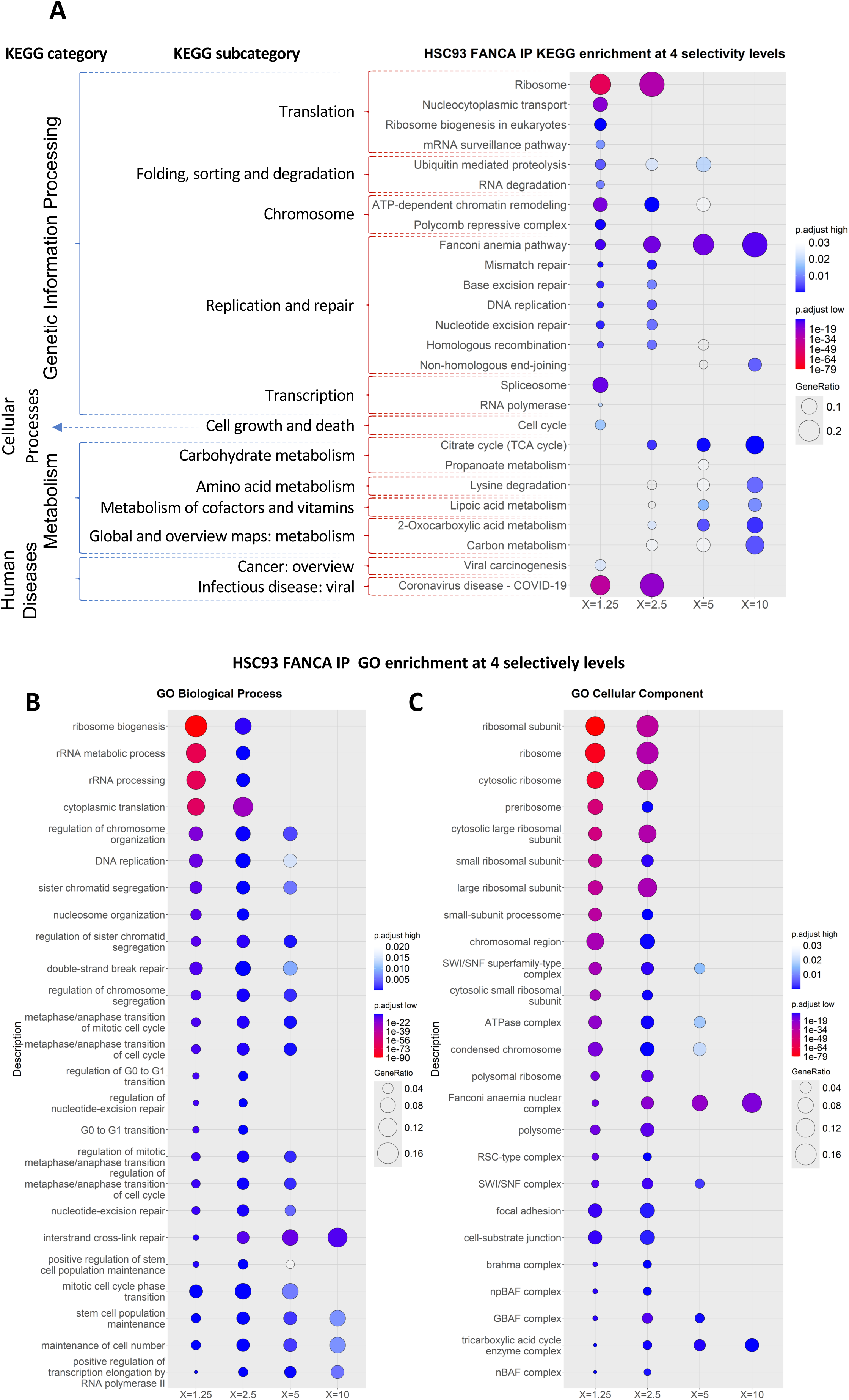
Functional enrichment analysis of FANCA co-IP preys at different level of selectivity. **(A)** Most significantly enriched functional KEGG**, (B)** GO_BP, and **(C)** GO_CC terms in HSC93 FANCA IP at levels **X**=1.25, **X**=2.5, **X**=5, **X**=10.

We next conducted functional analyses on the partners identified by IP in RPE1 cells and by BioID in Flp-in HEK293 cells. We used the publicly available web platform STRING (v. 12.0) to perform functional enrichment analysis (44). The number of enriched KEGG functional terms found by STRING with the HSC93 IP datasets **X**=5, **X**=10, SP>0.75 and SP>0.95, was too low to perform any meaningful comparison. Therefore, we submitted to STRING the protein lists from selectivity levels **X**=1.25 and **X**=2.5, identifying 14 and 9 functional enriched KEGG terms (FDR<0.5; strength>0.5), respectively (Figure S2; Table ST6). We performed the same analysis on the RPE1 IP-MS data and HEK293 BioID at IR≥1.5, and IR≥5. First, we noticed that the KEGG terms retrieved by STRING from the **X**=1.25 and **X**=2.5 lists were generally, although not entirely, replicating the results previously found with ClusterProfiler (Figure 5A *vs* 4A; Figure S2), reflecting differences between STRING and ClusterProfiler, possibly due to different *p*-value correction methods, since both algorithms rely on the same hypergeometric test method. KEGG term enrichment by STRING in the RPE1 IP (Figure 5B; Figure S2) showed terms related to: translation (“Ribosome” being the most significantly enriched term), mRNA processing (“Spliceosome”, “mRNA surveillance pathway”, “RNA transport”), “Fanconi anemia pathway”, and, mitochondria (“TCA cycle” and “Mitophagy”, a process described as altered in FA (26)), in addition to several terms associated with different DNA transactions (Figure 5B). Finally, KEGG enrichment analysis applied to both BioID lists recovered “Fanconi anemia pathway”, “Spliceosome”, “Homologous Recombination” and “RNA transport” at IR≥5 plus “Ribosome”, “MicroRNAs in cancer” and “Amyotrophic lateral sclerosis” (but not “Homologous Recombination”) at IR≥1.5 (Figure 5C; Figure S2; Table ST6, Sheet 1).

**Figure 5.**
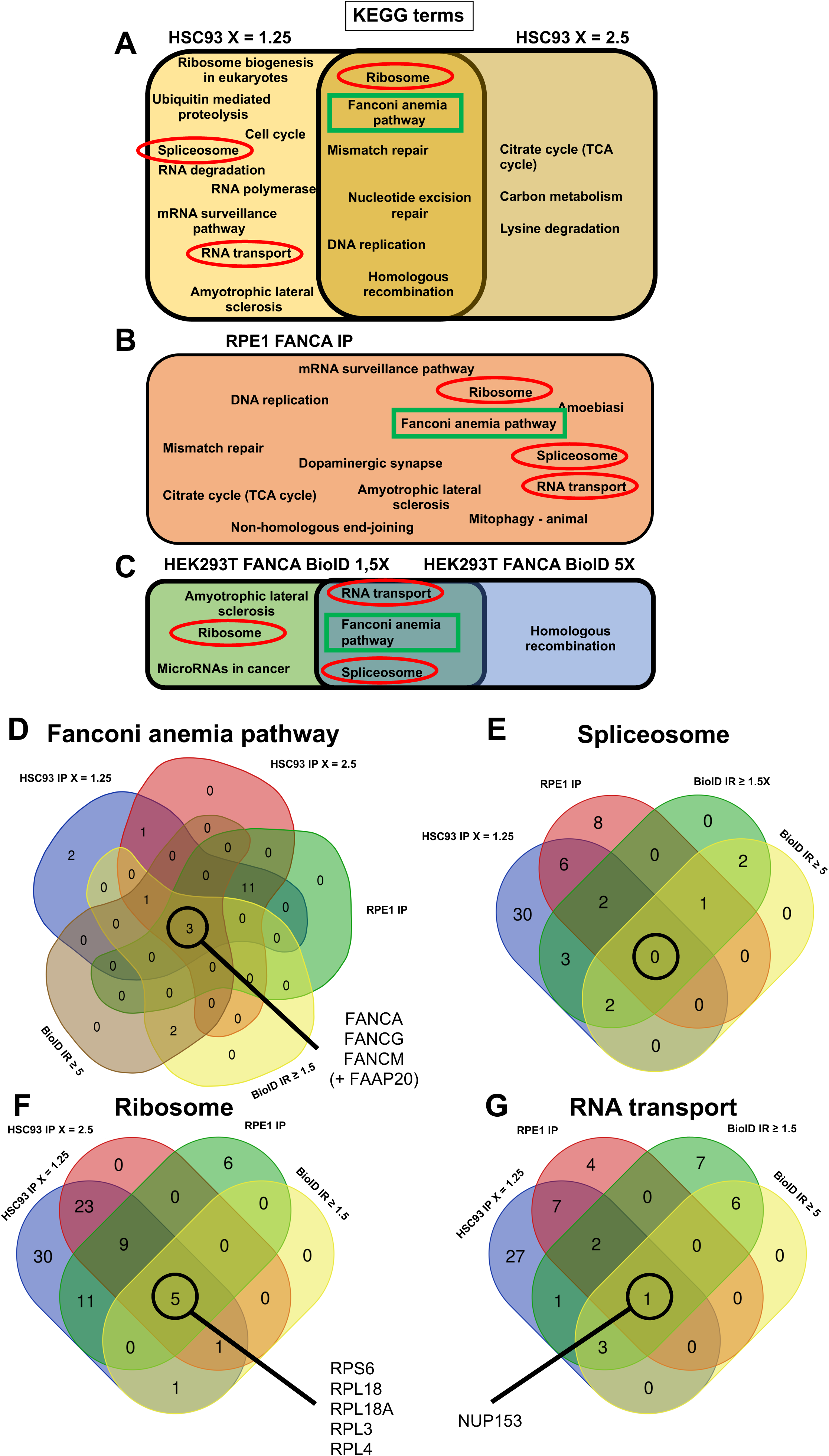
Functional enrichment analysis of FANCA preys from different datasets. **(A)** Venn diagram representing the overlap of enriched KEGG terms between the **X**=1.25 and **X**=2.5 datasets from the co-IP in HSC93 cells. **(B)** KEGG enriched terms emerging from the FANCA partners identified in the RPE1 cells. **(C)** KEGG enriched terms identified in the BioID>1.5 and BioID>5 lists issues from the Flp-in HEK293 cell expressing the FANCA fused with BirA. **(D to G)** Venn diagram representing the overlap of genes contributing to the enrichment of KEGG term “Fanconi anemia” **(D)**, “Spliceosome” **(E)**, “Ribosome’ **(F)**, and RNA transport **(G)** in the above reported datasets we generated.

Next, we compared the KEGG terms enriched in the five aforementioned protein lists, as well as the proteins specifically responsible for their enrichment. Only one term, “Fanconi anemia pathway” (circled in green), was found enriched across all five datasets, and 3 terms, “Ribosome”, “RNA transport” and “Spliceosome” (circled in red), were enriched in four out of five (Figure 5A-C). Overall, different experimental approaches (IP vs Bio-ID), in different cell models (HSC93, RPE1, HEK293) allow identifying similar terms. We next examined whether the proteins participating to common enriched terms across different protein lists are the same across datasets (Figure 5D to G). The term “Fanconi anemia pathway” emerged thanks to presence of 18, 16, 14, 6 and 6 proteins in the five lists (**X**=1.25, **X**=2.5, RPE1, BioID IR≥1.5 and IR≥5, respectively). Strikingly, only three proteins are common to the five lists: FANCA, FANCG and FANCM (Figure 5D). Curiously, the protein FAAP20, the third component of the FANCA-FANCG-FAAP20 subcomplex, and whose loss-of-function in mice (no human patients reported to date) is known to induce a mild FA-like phenotype with reproductive and hematopoietic defects as well as MMC hypersensitivity (45), is not annotated in the KEGG registry as part of the term “Fanconi anemia pathway”. The term “Ribosome” was enriched due to the presence of 80, 38, 31, and 20 proteins in the HSC93 IP **X**=1.25, RPE1 IP and the two BioID lists, respectively. Nevertheless, only 5 proteins, RPS6, RPL18, RPL18A, RPL3, and RPL4, are in common among the four datasets, and, notably, all but one (RPS6), are constituents of the 60S large ribosomal sub-unit. Finally, the term “RNA transport” shows only one common protein across datasets (NUP153, a nuclear pore complex component) whereas no protein is shared by all datasets under the term “Spliceosome” (Figure 5E to 5G).

The observation of few, if any, common proteins contributing to the enrichment of common functional term across independent datasets highlights both the capability and necessity of using diverse approaches to identify common biological pathways and functions affected by a single protein through its PPI and proxilome landscapes. We also analyzed the list of FANCA preys identified by Lagundžin et al. (31) and that compiled in the BioGRID database (Table ST3, Sheet 2). In the first list (139 proteins), only 4 enriched KEGG terms, all related to DNA transactions, emerged (Table ST3, Sheet 2). In the second (159 proteins), the STRING functional analysis retrieved as many as 77 KEGG enriched terms (FDR>0.05; strenght >0.5) (Table ST3, Sheet 2). The large majority of the 77 enriched terms, *i.e.* 65, referring to cancer and cell signaling, emerges as a consequence of the presence of a small set of proteins comprising KRAS, NFKB/IKBKB, AKT1 and NOTCH1. Apart from two terms (’MicroRNAs in cancer’ and’Amyotrophic lateral sclerosis’), all the other refer again to DNA processes (Table ST3, Sheet 2).

In conclusion, our in silico analysis supports the involvement of FANCA in several distinct networks: one, expected, related to DNA repair and chromatin/chromosome behavior, a second one involved in RNA processing, a third one involved in nucleocytoplasmic trafficking, a fourth one involved in translation and, finally, a fifth one involved in several metabolic aspects, unveiling a key position of FANCA in the cellular physiology.

### A multilayered in silico FANCA PPI landscape

Using complementary experimental and analytical approaches, we generated multiple FANCA interaction datasets. We subsequently validated our findings against published data, confirming that our datasets link FANCA not only to its canonical roles in DDR but also to other biological functions. The contribution to these functions may include direct biochemical/physical interactions, indirect connections mediated by other proteins, and participation to common subcellular structures. To delineate the FANCA PPI network we used the 9 protein-coding gene lists we generated (Table ST3, Sheet 1 without the list of proteins common in **X**=10 and SP>0.95) together with the list of the 316 FANCA interactors found in literature to create five additional catalogues (AC) of FANCA partners corresponding to proteins present, respectively, in all, at least 9, 6, 3 and 2 of the original gene lists (AC_10 to AC_2). ACs are not mutually exclusive, *i.e.* AC_2 includes the proteins in AC_3, which includes those in AC_6, and so on. Thus, AC_10 contains 3 proteins, AC_9 5 (3+2), AC_6 56 (51+5), AC_3 235 (179+56), AC_2 500 (265+235) (Table ST3, Sheet 5). Our aim is to generate a multi-layered landscape of FANCA partners, each layer corresponding to a degree of strength and/or proximity and/or abundance of the interaction with FANCA.

We used STRING to identify the enriched KEGG, GO_BP and GO_CC terms in AC_2, the least selective of ACs, and obtained (FDR<0.05, strength >0.5) 11 KEGG, 220 GO_BP and 90 GO_CC terms (Figure S3). However, the association between proteins and functional KEGG or GO terms is subject to potential annotation bias from two main sources: a) differences in the protein sets defining a given term between KEGG and GO databases, and, b) the fact that different terms often refer to biologically related events/pathways. For instance, the term “Ribosome” gathers 131 proteins in KEGG (including 50 in AC_2) and 228 GO_CC (58 of which are in or AC_2). In AC_2, respectively, 51, 65, and 36 proteins are associated to GO_BP terms “Cytoplasmic translation” (containing 123 proteins), “Translation” (389 proteins), and “Regulation of translation” (456 proteins). This results in 85 unique translation-related proteins in FANCA’s PPI landscape.

We assembled the proteins responsible for the emergence of 7 key and closely related enriched terms from the KEGG and GO databases in the AC_2 list and assessed their presence in the most stringent ACs. This strategy allows us to propose a multi-layered FANCA PPI landscape (Figure 6; Table ST3, Sheet 6-12).

**Figure 6.**
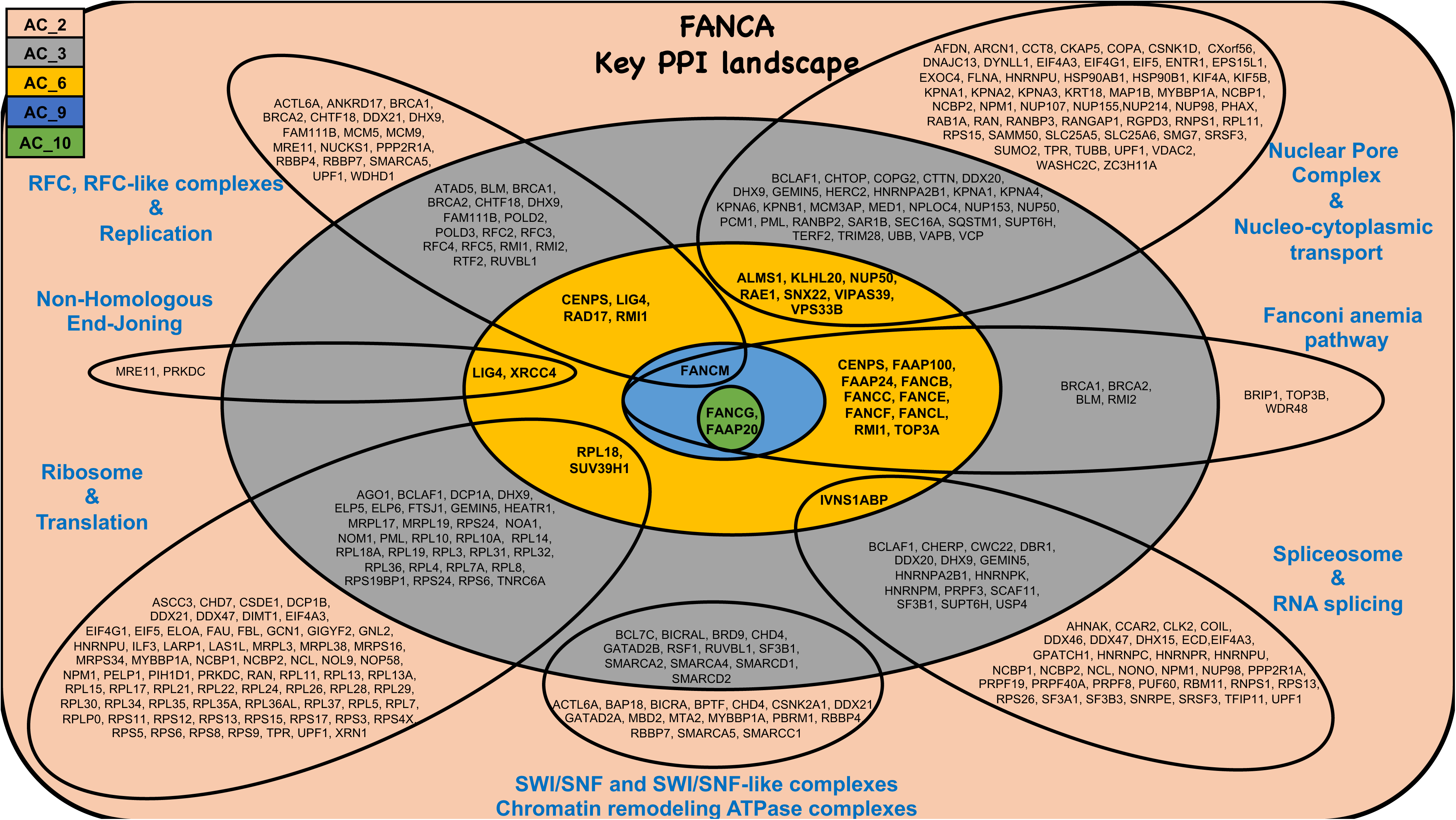
FANCA PPI landscape. Schematic representation of the major sub-networks that characterize the FANCA interactions landscape.

Associating the proteins belonging to the KEGG term’Fanconi anemia’ (51 proteins) with GO term “Fanconi anemia nuclear complex” (14 proteins), we identified 21 unique proteins in AC_2 (corresponding to 20/51 and 12/14, respectively, in the KEGG and GO_CC lists). Three of them, the components of the AG20 sub-complex (FANCA-FANCG-FAAP20), are the only proteins present in the most selective layer, *i.e.* AC_10, whereas 4, 14 and 18 were in the AC_9, _6, and _3, respectively. According to the representation we propose (Figure 6), the connection between FANCA and proteins involved in processes distinct from genome integrity maintenance shows a level of proximity comparable to that between FANCA and most members of the FANC core complex. Examples include proteins related to nucleocytoplasmic trafficking (NUP50, RAE1, SNX2), RFC-like complexes (RAD17), large ribosome subunit (RPL18), mRNA splicing (IVNS1ABP), and NHEJ (LIG4, XRCC4).

We postulate that FANCA plays a role in ribosome biogenesis, mRNA splicing, NHEJ, and nucleocytoplasmic trafficking through interactions with some of their key players, such as RPL18, IVNS1ABP, LIG4-XRCC4 and NUP50/RAE1.

### The FANCA PPI network strengthens its biological connection to ribosome biogenesis/translation

To biologically validate our findings, we reiterated FANCA IP in extracts from HSC93, HeLa, Hela-Kyoto, U2OS, HEK293 and RPE1 cell lines under varying conditions (150 nM vs. 300 nM salt; with or without benzonase; exposure to Mitomycin C or Aphidicolin). This time the IP was followed by immunoblotting. We successfully recovered previously reported interacting proteins of FANCA, *i.e.* FANCG, RPL18, NPM1, and NCL as well as undescribed interactors that we had identified by IP-MS, *i.e.* CUX1, IVNS1ABP, RAE1, LIG4, and XRCC4 (Figure 7A to G and 7H). We demonstrated that the interaction between FANCA and FANCG is resistant to both salt concentration and benzonase treatment, whereas the interaction with NCL and NPM1, is lost at high salt concentration and diminished, but still undoubtedly detected, after benzonase treatment as is the case for RPL18 (Figure 7A and 7B). These data are consistent with a direct physical FANCA-FANCG interaction and suggest a contribution of RNA in the association of FANCA with other partners of its PPI landscape, supporting a participation of FANCA in supra-molecular assemblies, known as condensates (46). The FANCA-CUX1, FANCA-RAE1 and FANCA-LIG4/XRCC4 co-IPs persist in response to MMC or APH in both human lymphoblasts, HEK293 and/or RPE1 cells (Figure 7D to 7F). Moreover, the level of MMC-induced chromatin association of LIG4 as well as that of other NHEJ components such as 53BP1 and KU70, is increased in FANCA-deficient cells (Figure 7H). However, we cannot conclude that this effect is specifically mediated by the loss of the FANCA-LIG4/XRCC4 interaction, since over-and/or unscheduled activation of the NHEJ pathway is a general characteristic of all FA cells. Finally, as a measure of specificity of our IP/WB, we compared the co-IP of NPM1, NCL, and RPL18 with FANCA or FANCD2 as baits. While NCL and NPM1 are also co-IP by FANCD2 but at a significantly lower level than FANCA, FANCD2 failed to co-IP RPL18 and FANCG, the key FANCA interactor (Figure 7I).

**Figure 7.**
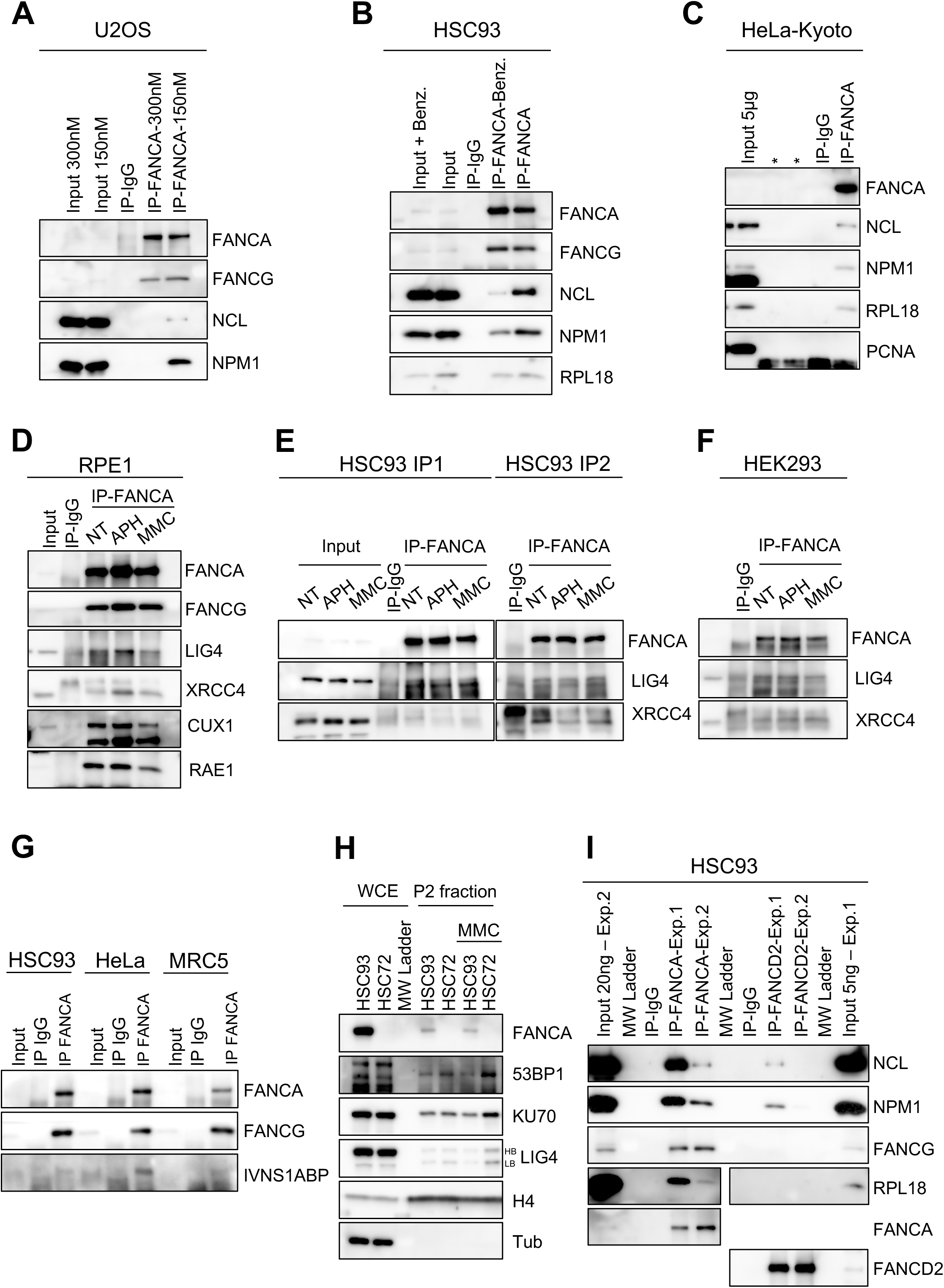
Validation of the FANCA co-IP. (A-C) FANCA co-IP of FANCG, NCL, NPM1 and RPL18 in U2OS, HSC93 or HeLa-Kyoto cells in unstressed conditions(A-C) under different salt (A) or benzonase (B) conditions. **(D-F)** FANCA co-IP of FANCG, LIG4 and XRCC4, CUX1 and RAE1, in unstressed conditions as well as in response to Mitomycin C (MMC, 200ng/ml, 16-24h) or aphidicolin (APH, 0,6µM, 16-24h) in RPE1 (D), HSC93 (E) or HEK2983 (F) cells. **(G)** FANCA and INVS1ABP co-IP in HSC93, HeLa or MRC5 cells. **(G)** Chromatin association of FANCA, 53BP1, KU70 and LIG4 in HSC93 and HSC72 cells in instressed conditions and in response to Mitomycin C (MMC, 200ng/ml, 16-24h). H4 and Tubulin (Tub) are used as control for chromatin extraction efficiency. **(H)** Co-IP of NCL, NPM1 FANCG, RPL18 with FANCA or FANCD2 in HSC93 cells. Two experiments with similar outcomes were reported.

Although characterizing the contribution of FANCA to the different protein networks identified is beyond the scope of this work, we addressed the question of the biological significance of the presence of many ribosomal proteins in FANCA’s PPI landscape. We performed polysome profiling (Figure 8A) and analyzed the distribution of proteins from the large (LSU) and small (SSU) ribosomal subunits (RPL and RPS proteins, respectively) in the 40S, 60S, 80S and polysome fractions (Figure 8 and Table ST7, Sheet 1 and 2). Beyond the already observed accumulation of monosomes (80S) in FANCA-deficient (HSC72) compared to FANCA-proficient cells (HSC93) as evidenced by the A254 profile (Figure 8A) (27), immunoblot analysis of the fractions showed an increase in RPL and RPS proteins in pre-polysome fractions (40S, 60S, and 80S) and a decrease in polysome fractions in FANCA-deficient cells compared to-proficient cells (Figure 8B to 8D). This result, indicating a decreased amount of actively translating ribosomes (*i.e.* polysomes) supports our previous observation that FANCA loss negatively affects the translation rate (27).

**Figure 8.**
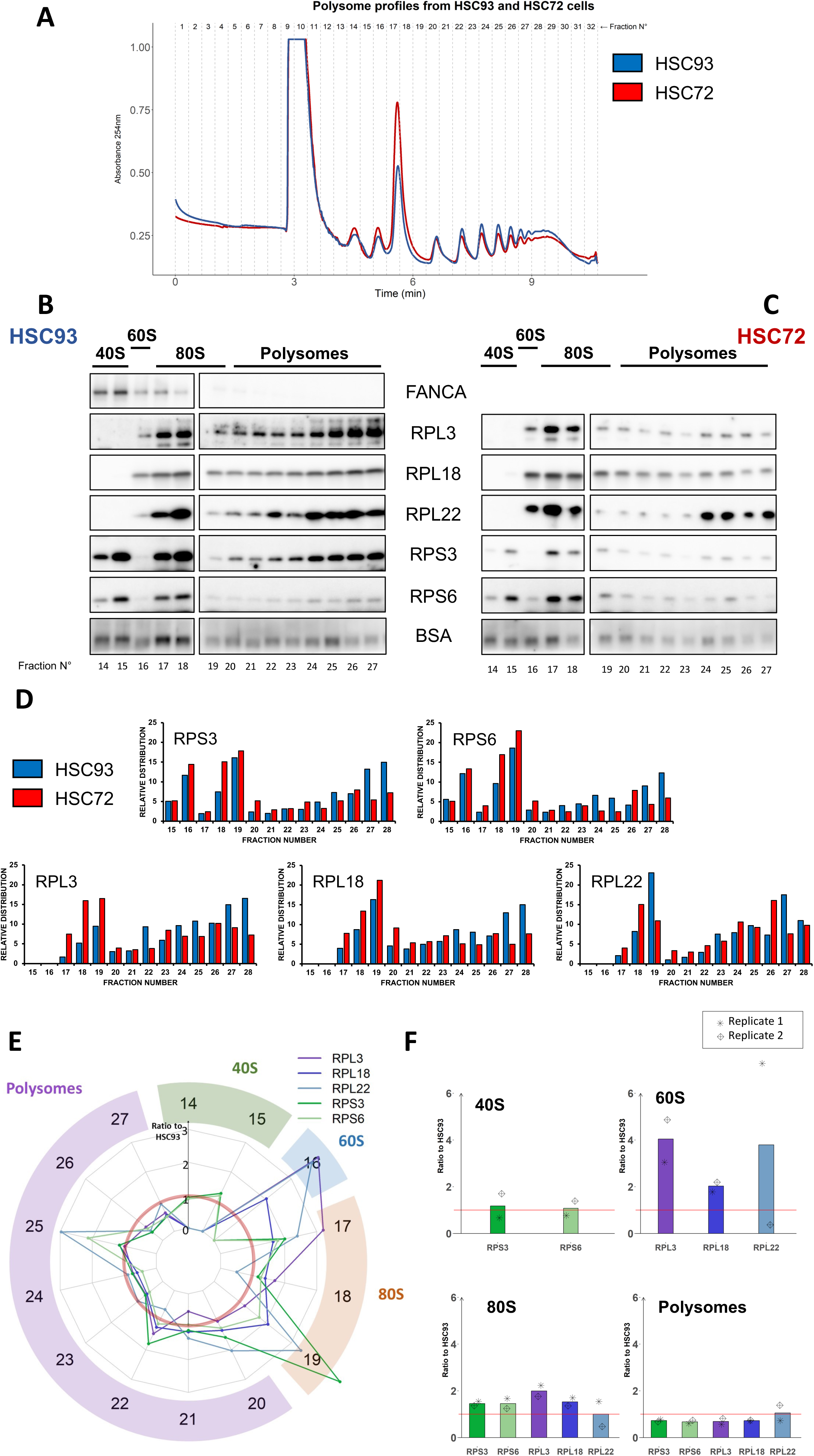
FANCA contributes to ribosome stoichiometry. **(A)** Polysome profiling in exponentially growing HSC93 and HSC72 cells. **(B-C)** Western blot showing the distribution of indicated RPS and RPL proteins in the 40S, 60S, 80S and polysomal fractions. BSA (dose) added in each fraction before TCA extraction from glycerol fraction was used as standard for quantification (Spike-in), in HSC93 and HSC72 cells. **(D)** Relative distribution of the indicated proteins in the indicated fractions in HSC93 and hSC72 cells. Data were from replicate 1. **(E)** Relative ratio of indicated proteins in the 40S, 60S, 80S and polysomal fraction in the HSC72 cells vs the FANCA WT HSC93 cells. Mean of two replicates. **(F)** Distribution of analyzed RPL and RPS proteins in the different ribosomal sub-units in the absence of FANCA. Red line indicates the level of the same proteins in the FANCA-proficient cells settled to 1.

We repeated the same experiment a second time, independently (Table ST7, Sheet 2). Using the quantitative data from immunoblot detection of ribosomal proteins in both experiments (Table ST7, Sheets 1-4), we calculated, for each protein, the average signal (normalized to BSA) ratio between HSC93 (FANCA-proficient) and HSC72 (FANCA-deficient) cells, in each fraction (Figure 8E). Again, in extracts from FANCA-deficient cells, we observed a general increase in RPL proteins in the 60S fraction (N°16), and in RPL and RPS proteins in the 80S fractions (N°17 to 19), while the level of SSU proteins in the 40S fraction appears independent of the FANCA status of the cells. Notably, by grouping the fraction signals by RNP complex (Figure 8F), it appears that the ratio for each protein is similar in each individual experiment for 40S, 80S and Polysomes, with all tested LSU and SSU proteins appearing increased in 80S and decreased in Polysomes, with the exception of RPL22 whose WT/FANCA ratio remains equal to one (Figure 8F). In 60S, the average WT/FANCA ratio for RPL3, RLP18 and RPL22 indicates a higher level of all three proteins in FANCA extracts. However, it can be noted that the variation between the two experiments in 60S is greater than in 40S, Monosomes and Polysomes, especially with RPL22, for which no tendency can be drawn. Our data suggest that FANCA deficiency alters LSU protein stoichiometry, particularly in the 60S complex. In other words, the assembly and final processing (i.e., its stoichiometry) of 60S subunit RPLs, as well as their association with the 40S to form a productive monosome, appear to be impaired in the absence of FANCA, thereby impairing translation. This hypothesis is consistent with the role of FANCA in ribosome biogenesis, since FANCA is present in the 40S, 60S, and 80S fractions (Figure 8B).

Overall, our observations support a functional contribution of FANCA to ribosome biogenesis/translation, likely through interacting with RPL components, including RPL18.

## Discussion

Our analysis provides an overview of the FANCA PPI landscape, confirming known partnerships and revealing novel associations that extend FANCA’s contribution to biological functions beyond the canonical FANC/BRCA pathway and DDR. While the current hypothesis postulates that the various phenotypic abnormalities associated with loss of the FANC/BRCA pathway are due to dysfunctions secondary to alterations in the cells’ DDR, our observations support an alternative explanation. FANCA’s connectivity landscape appears to intersect a broader range of processes than previously suspected. Indeed, beyond its known partners within the FANC/BRCA pathway, FANCA appears associated with proteins and complexes involved in NHEJ, chromatin dynamics, nucleocytoplasmic transport, RNA metabolism, ribosome biogenesis, and translation. These results were made possible by the analysis of low-selectivity FANCA partners, and the combination of results gathered from several distinct approaches This strategy made possible the detection of an enrichment in functional terms of which only a few representatives are present in the higher-selectivity datasets, which would therefore have been missed by only using the canonical SAINT algorithm.

Regarding the DDR and genetic stability maintenance pathways, we retrieved the FANCcore complex along with FANCM and its partners, and extended the already reported biochemical and/or functional connections of FANCA or the FANC/BRCA pathway with several complexes of the RFC-like family (Figure S4A), whose activity contributes to the loading/unloading of PCNA and 9-1-1 complexes onto the chromatin (47, 48) for both DNA replication and repair. Moreover, our data extend previous observations linking FANCA to the NPC and other proteins involved in transport across the nuclear membrane that contribute to DNA repair (Figure S4B). Notably, it has been described that NUP153, NUP50 and the nuclear pore-associated SUMO protease SENP1 contribute to the nuclear transport of the NHEJ-associated protein 53BP1 (49) fostering its assembling in subnuclear foci at DSB sites (50–52). The known unscheduled usage of NHEJ over HR in FANC/BRCA pathway-deficient cells has been associated with an accumulation of 53BP1 foci. In our FANCA IP, we identified NUP153, NUP50 and SENP1 as well as the XRCC4-LIG4 heterodimer and DNA-PKcs and validated the known increase in chromatin association of 53BP1, KU70 and LIG4 in FANCA-deficient cells treated with MMC (53, 54). FANCA could therefore influence the NHEJ/HR balance through two new mechanisms: (a) regulating 53BP1 sorting to chromatin by interacting with NUP153-NUP50-SENP1, and (b) interacting with the XRCC4-LIG4, thereby controlling its activity. These two actions on the NHEJ/HR balance would add to the already known ones: supporting HR by acting as a backup for RAD52 during SDSA (55) and BRCA2/RAD51-dependent HR rescue through FANCD2/FANCI (Figure S4C).

We also extended the potential functional scope of FANCA in chromosome maintenance and dynamics by identifying its association with several chromatin remodeling complexes, including the INO80-type, ISWI-type, NuRF-type, NuRD-type, and the mammalian SWI/SNF superfamily (Figure S5). These complexes play essential roles in maintaining genome stability (Figure S4C) and regulating RNA expression. In particular, beyond SMARCA4 (also known as BRG1), a previously established FANCA interactor (34), whose depletion recapitulates multiple Fanconi anemia-like cellular phenotypes (e.g., reduced EdU incorporation, increased replication fork asymmetry, enhanced R-loop formation and associated γH2AX foci, elevated micronuclei and ana-telophase bridges, disrupted nucleolar architecture, increased sensitivity to PARP inhibitors and cisplatin, and impaired RAD51 loading) (56–59), we also identified BRD9, BICRA, and BICRAL, three unique components of the non-canonical BAF (ncBAF) complex. Notably, BRD9 depletion has been linked to reduced rRNA transcription, lower mRNA association with polysomes, and globally decreased translation rates (60), phenotypes closely mirroring those we recently observed in FANCA-deficient cells (27). Furthermore, BRD9 plays a critical role in hematopoietic cell identity and lineage commitment. Its loss has been shown to support colony-forming unit (CFU) activity, promote myeloid skewing, reduce stemness, increase the fraction of actively cycling LSK cells with elevated γH2AX levels, and impair erythroid differentiation—all of which are consistent with the hematopoietic defects observed in FA patients and mouse models (61). Altogether, these findings suggest that disruption of the FANCA–SWI/SNF axis may contribute significantly to both cellular dysfunction and clinical heterogeneity in Fanconi anemia.

Confirming our previous observations (27), FANCA associates with the 40S and 60S free ribosomal subunits and the 80S ribosomes, but not with polysomes (*i.e.* actively translating mature ribosomes), supporting the involvement of FANCA in the process of ribosome assembling, possibly mediated by its interaction with RPL18 previously identified in a yeast two-hybrid screen (32) and that we validated here. Indeed, RPL18 happens to be the only cytoplasmic ribosomal protein passing the in-house filter at selectivity level **X**=5. Notably, supporting a functional meaning of the FANCA-RPL18 or the FANCA-ribosome partnership, FANCA loss alters the ribosomal protein stoichiometry mainly inside the 60S subunit. Such alterations could contribute to the reduced rate of translation we reported in FANCA-and FANCG-deficient cells.

Finally, FANCA appears connected to several partners involved in mRNA metabolism, and future works will determine how FANCA impacts mRNA transcription, splicing, transport and/or stability. Intriguingly, it was demonstrated that FANCA binds to RNA with an affinity much greater than that measured for single-stranded (ssDNA) and double-stranded (dsDNA) DNA (*K_d_* of 2.8, 11.1 and 42.5 nM, respectively) (62). Most of the potential FANCA partners we identified are ribonucleoproteins or complexes able to associate through weak intermolecular bonds forming molecular sub-cellular membrane-less structures or condensates, that enhance intermolecular and enzymatic interactions, fostering the efficiency and accuracy of the biological processes (63). The presence of RNA molecules appears mandatory for the aggregation of the majority, if not the totality, of such condensates. FANCA, as also FANCI, loss has been associated with abnormalities in at least two of such condensates: the nucleolus (27) and nuclear speckles (64). Therefore, even if the PPI landscape of FANCA we present here could be influenced by its affinity for RNA, its potential participation to several subcellular condensates could contribute to their’stability’ influencing multiples aspects of the cellular physiology. Beyond RNA, the presence of proteins with intrinsically disordered regions (IDRs) is another main determinant of biomolecular condensation. Evaluated thanks to several predictive algorithms (65–67), FANCA presents two short regions, in the 3’ and 5’ tails of the protein presenting IDR sequences exceeding the threshold value of 0.5 (Figure S6A to S6C). Strikingly, its key partner FAAP20 is predicted to be practically entirely (90%) disordered (Figure S6). Notably, 9% and 43% of the disordered residues in FANCA and FAAP20, respectively, are predicted be RNA-binding (Figure S6C), supporting the possibility that the potential participation of FANCA to biomolecular condensates is mediated by both its affinity for RNA and its association with FAAP20. Such FANCA behavior may underlie the pleiotropic and heterogeneous nature of the syndrome.

In conclusion, beyond the field of FA, our approach underscores the importance of reevaluating the emphasis on the “strongest” interactions or other prominent biological links between cellular components as the central focus for describing cellular physiology. Indeed, information derived from the integrative analysis of “weaker” interactions and the combination of several alternative approaches may be more meaningful to decipher the various interconnected networks and pathways contributing to cell physiology. Studying each distinct pathway and process associated with FANCA and integrating them into a comprehensive model is beyond the possibilities of a single article. However, our data pave the way for further analyses to validate and expand the significance of the FANCA-PPI network. This could open new therapeutic avenues for patients and enable anticipating and counteracting the two most severe manifestations of Fanconi anemia, *i.e.* BMF and oncogenesis.

## Materials and Methods

### Cell lines, culture conditions and treatments

EBV-immortalized lymphoblastoid cell lines HSC93 (FANCA wild type) and HSC72 (FANCA deficient), were grown in RPMI supplemented with 13% fetal bovine serum, 100 U/ml penicillin and 100 µg/ml streptomycin (all from Invitrogen). HeLa, HeLa-Kyoto, U2OS, HEK293, and MRC5 were grown in Dulbecco’s modified Eagle’s medium (DMEM) (Life Technologies) supplemented with 10% fetal calf serum (FCS), penicillin (0.5 mg/ml), streptomycin (100 μg/ml), and 1 mM pyruvate. RPE1-hTERT cells were grown in DMEM-F12 supplemented with 10% FBS and penicillin-streptomycin.

Flp-In 293 T-REx and the Flp-In 293 T-REx derived stable cell lines were grown under standard sterile cell culture conditions in Dulbecco’s modified Eagle’s medium (DMEM, Merck-Sigma-Aldrich, D5796) containing 10% fetal bovine serum (BioWest S1810-500) and penicillin-streptomycin. Parental cells were selected with 100 µg/ml Zeocin (ThermoFisher Scientific R25001) and 15 µg/ml Blasticidin (InvivoGen, ant-bl) and Flp-In 293 T-REx derived stable cell lines were maintained with 5 µg/ml Blasticidin (InvivoGen, ant-bl) and 50 µg/ml Hygromycin B (Sigma-Aldrich, H3274).

All cells were cultivated at 37°C under a 5% CO2 atmosphere and were routinely tested for mycoplasma and scored negative.

Mitomycin C (Sigma-Aldrich, M0503) and aphidicolin (Sigma-Aldrich, A0781) were prepared in H_2_O and added for 18 to 24 hours.

### Protein extraction and immunoprecipitation

Cellular extracts for immunoprecipitation were prepared from exponentially growing lymphoblasts or adherent cells. Cells were lysed on ice for 30 min in NaCl, EDTA, Tris, NP-40 (NETN) buffer (150 mM/300 mM NaCl, 50 mM tris (pH 8.0), 0,5% NP-40, 1 mM EDTA), supplemented with phosphatase inhibitor PhosSTOP (Roche), cOmplete ULTRA EDTA-Free Protease inhibitors (Roche), benzonase (Merck, 1 mM) and 1 mM MgCl2. Extracts were sonicated for 2 × 10 s at 30% (Vibracell 75042, Bioblock) and spun down for 5 min at 13,000 rpm. Supernatants were quantified using the Bradford assay (BioPhotometer, Eppendorf). One milligram of protein extract was used per immunoprecipitation with rabbit anti-FANCA were from Bethyl (A301-980); rabbit anti-IgG were obtained from Dako.

For each immunoprecipitation, antibodies (3 μg) were coupled to 20 μl of magnetic beads (Dynabeads Protein G Magnetic Beads, Thermo Fisher Scientific). Beads washed in 1× PBS (two times) were resuspended in 300 μl of 1× PBS and incubated with antibodies for at least 2 hours in a roll shaker in a cold room. To preclear the protein extracts, 10 μl of beads washed in PBS and in NETN buffer was added to each whole protein extract in the presence of 0.5 μg of control immunoglobulin G (IgG) and incubated for at least 2 hours in a roll shaker in a cold room. Beads coupled with IgG were collected on a magnetic support and discarded. Supernatants were mixed with beads coupled to specific antibodies. A small amount of the precleared protein extract was kept as input. Precleared supernatants were incubated with the magnetic beads coupled to the antibodies overnight in a roll shaker in a cold room. Beads coupled with antibodies were captured on a magnetic support and washed four times with 20µL-40µl of NETN buffer before being resuspended in 40 μl of 2× Laemmli and heated for 5 min at 70°C to dissociate proteins and antibodies from the beads. The supernatants were transferred into a new tube, 0,5-1 μl of β-mercaptoethanol was added. For WB analysis, immunoprecipitated proteins are loaded on an acrylamide gel after 5 min of heating at 98°C.

### Proteomics analysis on IP samples Sample preparation

The Co-IP samples were solubilized in lysis buffer (2% SDS, 200mM Tris-HCl, pH 8.0, 10 mM TCEP, 50 mM chloroacetamide). Bottom-up experiments’ tryptic peptides were obtained by Strap Micro Spin Column according to the manufacturer’s protocol (Protifi, NY, USA). Briefly: Proteins were digested during 14h at 37°C with 1µg Trypsin sequencing grade (Promega). The Strap Micro Spin Column was used according to the manufacturer’s protocol. After speed-vacuum drying, eluted peptides were solubilized in 2% trifluoroacetic acid (TFA).

### Liquid Chromatography-coupled Mass spectrometry analysis (nLC-MS/MS)

nLC-MS/MS analyses were performed on a Dionex U3000 HPLC nanoflow chromatographic system (Thermo Fischer Scientific, Les Ulis, France) coupled to a TIMS-TOF Pro mass spectrometer (Bruker Daltonik GmbH, Bremen, Germany). Peptides were solubilized in 10µl of 0.1% trifluoroacetic acid (TFA) in 10% Acetonitrile (ACN). One μL was loaded, concentrated and washed for 3min on a C18 reverse phase Pepmap neo (3μm particle size, 300 μm inner diameter, 5 mm length, from Thermo Fisher Scientific). Peptides were then separated at 50°C on an Aurora C18 reverse phase resin (1.6 μm particle size, 100Å pore size, 75μm inner diameter, 25cm length) (IonOpticks, Middle Camberwell Australia) with a 60 minutes overall run-time gradient from 99% of solvent A containing 0.1% formic acid in milliQ-grade H2O to 40% of solvent B containing 80% acetonitrile, 0.085% formic acid in mQH2O with a flow rate of 400 nL/min. The mass spectrometer acquired data throughout the elution process in a positive mode and operated in DIA PASEF mode with a 1.38 second/cycle, with Timed Ion Mobility Spectrometry (TIMS) enabled. Capillary voltage was set to 1,500 V. Ion accumulation and ramp time in the dual TIMS analyzer were set to 100 ms each. The MS1 spectra were collected in the m/z range of 100–1,700. The diaPASEF window scheme ranged in dimensions from m/z 400 to 1,200 and in dimension 1/K0 from 0.63–1.43. The collision energy was set by linear interpolation between 59 eV at an inverse reduced mobility (1/K0) of 1.60 versus/cm2 and 20 eV at 0.6 versus/cm^2^.

### Protein identifications and quantifications

The mass spectrometry data were analyzed using DIA-NN version 1.8.1 (68). The database used for *in silico* generation of spectral library was a concatenation of Human sequences from the Swissprot database (release 2022-05) and a list of contaminant sequences from Maxquant and from the cRAP (common Repository of Adventitious Proteins). M-Terminus exclusion and carbamidomethylation of cysteins was set as permanent modification and one trypsin misscleavage was allowed. Precursor false discovery rate (FDR) was kept below 1%. The “match between runs” (MBR) and the normalization option was allowed.

The results files of DIA-NN were analysed using a R script homemade. T-test greater Unpaired were done on proteins showing at least 3 valid values in one group and at least 70% of valid values in the other group using log2(LFQ intensity). Significant threshold is PValue<0.05.

### BioID assay

A doxycycline-inducible cDNA encoding FANCA fused to the mutant biotin ligase BirA* and a FLAG epitope was stably integrated into Flp-In HEK293 cells. The R118G mutation in BirA (BirA*) enables the transfer of biotin to nearby proteins, which are subsequently enriched using streptavidin-coated beads and identified by MS.

### Plasmid construct

To generate pDEST-pcDNA5-BirA-FLAG N-term-FANCA, the full-length FANCA cDNA from the pDONR-FANCA construct was inserted into the pDEST-pcDNA5-BirA-FLAG N-term vector (Gift from Anne-Claude Gingras) using a Gateway recombination assay.

### Generation of stable cell lines

Flp-In 293 T-REx cells are seeded to reach 80-90% confluence on the day of transfection. pDest-pcDNA5-BirA-Flag-FANCA expression plasmid was mixed with pOG44 encoding the Flp recombinase (ThermoFisher Scientific, V600520) at a 1:7 ratio in opti-MEM (Gibco, 31985-047). For a single transfection in a 6 well plate, 500ng of the expression plasmid was mixed with 3.5mg of pOG44 in 250uL opti-MEM. Additionally, 8uL Lipofectamine 2000 Transfection Reagent (ThermoFisher Scientific, 11668-019) was added to 250uL opti-MEM. After an incubation period of 5 minutes at room temperature, both solutions were mixed and incubated for a further 15 minutes at room temperature. The mixture was then pipetted dropwise onto the cells. The medium was changed after 6 hours. At 48 hours post-transfection, the cells were transferred to a 100mm petri dish, and 24 hours later, the selection was performed by adding 5 µg/mL Blasticidin and 50 µg/mL Hygromycin B. Clones were pooled, and the cells were examined for the expression of the construct by immunoblotting.

### Affinity capture of biotinylated proteins: BioID

Flp-In^TM^ 293 T-Rex cell lines stably transfected with BirA*-Flag-FANCA grown to 75% confluence were incubated with 1 µg/ml of doxycycline (Clontech, 631311) for 16h and with 50 mM biotin for 16 hours. Cells were washed with PBS and lysed with lysis buffer (50mM Tris-HCl pH 7.5, 150mM NaCl, 1mM EDTA, 1mM EGTA, 1% NP-40, 0.2% SDS, 0.5% Sodium deoxycholate) supplemented with 1X complete protease inhibitor (Roche, 4693159001) and 250U benzonase (Sigma, CE1014). Lysed cells were incubated on a rotating wheel for 1h at 4°C prior sonication on ice (40% amplitude, 3 cycles 10sec sonication-2sec resting). After 30min centrifugation (7750 rcf.) at 4°C, the cleared supernatant was transferred to a new tube and total protein concentration was determined by Bradford protein assay (BioRad, C500-0205). For each condition, 300 mg of proteins were incubated with 30ml of Streptavidin-Agarose beads (Sigma, CS1638) on a rotating wheel at 4°C for 3hr. After 1min centrifugation (400 rcf.), beads were washed, successively, with 1ml of lysis buffer, 1ml wash buffer 1 (2% SDS in H2O), 1ml wash buffer 2 (0.2% sodium deoxycholate, 1% Triton X-100, 500mM NaCl, 1mM EDTA, and 50mM HEPES pH 7.5), 1ml wash buffer 3 (250mM LiCl, 0.5% NP-40, 0.5% sodium deoxycholate, 1mM EDTA, 500mM NaCl and 10mM Tris pH 8) and 1ml wash buffer 4 (50mM Tris pH 7.5 and 50mM NaCl). Bound proteins were eluted from the agarose beads using 40 µl of 2X Laemmli Sample buffer and sent for mass spectrometry analysis

### Mass spectrometry

Sample digestion was essentially performed as described (Shevchenko et al., 2006). Briefly, proteins were loaded on a SDS-PAGE (BioRad, 456-1034) and, after short migration, a single band was excised. Proteins in the excised band were digested with Trypsin (Promega). The resulting peptides were analyzed online by nano-flow HPLC-nanoelectrospray ionization using a Qexactive HFX mass spectrometer (Thermo Fisher Scientific) coupled to a nano-LC system (Thermo Fisher Scientific, U3000-RSLC). Desalting and preconcentration of samples were performed online on a Pepmap® precolumn (0.3 3 10mm; Fisher Scientific, 164568). A gradient consisting of 0% to 40% B in A (A: 0.1% formic acid (Fisher Scientific, A117), 6% acetonitrile (Fisher Scientific, A955), in H2O (Fisher Scientific, W6), and B: 0.1% formic acid in 80% acetonitrile) for 120 min at 300nl/min was used to elute peptides from the capillary reverse-phase column (0.075 3 250mm, Pepmap®, Fisher Scientific, 164941). Data were acquired using the Xcalibur software (version 4.0). A cycle of one full-scan mass spectrum (375–1,500 m/z) at a resolution of 60000 (at 200 m/z) followed by 12 data-dependent MS/MS spectra (at a resolution of 30000, isolation window 1.2 m/z) was repeated continuously throughout the nanoLC separation. Raw data analysis was performed using the MaxQuant software (version 1.6.10.43) with standard settings. Used database consist of Human entries from Uniprot (reference proteome UniProt 2021_01) and 250 contaminants (MaxQuant contaminant database).

### Protein extraction and cellular fractionation and Western blot analysis

For proteins expression, cells collected by centrifugation or on Petri dishes were disrupted in lysis buffer [50 mM Tris-HCl pH 7.5, 20 mM NaCl, 1 mM MgCl2, 0.1% SDS and 1% benzonase (Novus), supplemented with protease and phosphatase inhibitors (Roche)]. After 20 min of incubation at room temperature, the protein concentration was determined using the Bradford assay, and samples were combined with 4× Laemmli buffer containing β-mercaptoethanol and denatured by boiling.

For cell fractionation, cell pellets were resuspended in solution A (HEPES pH 7.9 10 mM, KCl 10mM, MgCl2 1.5 mM, sucrose 12%, glycerol 10%, DTT 1 mM, protease and phosphatase inhibitors) supplemented with 0.1% Triton X-100. The samples were incubated for 5 min on ice. The cell pellets were centrifuged for 4 min at 1300 g, and the soluble proteins (S1) were eliminated. The nuclei were incubated in solution B (3 mM EDTA, 0.2 mM EGTA, 1 mM DTT, and protease and phosphatase inhibitors) for 10 min and then centrifuged for 4 min at 1700 g, and the soluble nuclear proteins (S2) were eliminated. After a wash in solution B, the chromatin (P2) was pulled by centrifugation at 16,1 g, resuspended in Laemmli 1X buffer containing β-mercaptoethanol and sonicated 2 × 10 s at 30% (Vibracell 75042, Bioblock).

For Western blot, proteins from immunoprecipitation, cellular extracts and cell fractions were separated by SDS‒PAGE by electrophoretic migration performed in 25 mM Tris, 192 mM glycine, and 0.1% SDS buffer. Semidry transfer was performed with a TransBlot cell apparatus (Bio-Rad) in transfer buffer composed of 2.5 mM tris-HCl (pH 8.3), 19.2 mM glycine, 0.01% SDS, and 20% isopropanol for 1 hour and 30 min at 20 V. Nitrocellulose membranes (Protran 0.2 mm, Amersham) were blocked for at least 1 hour in 0.1% PBS–Tween 20 and 5% milk and incubated with primary antibodies in PBS/0.1% Tween 20/5% milk. Visualization was performed using ECL (Life), Western Bright ECL or Quantum (Advansta), or West Femto ECL (Thermo Fisher Scientific) developer. Images were acquired using a CCD camera (General Electric) or Amersham Imager 600 (GE Healthcare). All western blot quantifications were performed using densitometry measures and ImageJ software.

Antibodies used were reported in Table ST8.

### Polysome profiling and fraction collection

Cycloheximide (100 μg/ml) was added for 5 min to cell cultures and maintained in PBS washes. Cells were fractionated in 5 mM tris (pH 7.5), 2.5 mM MgCl2, and 1.5 mM KCl buffer supplemented with cOmplete ULTRA EDTA-free protease inhibitors (Roche), cycloheximide (100 μg/ml), RNase inhibitor (Promega RNasin 0.2 U/μl), 2 mM DTT, 0.5% Triton, and 0.5% sodium deoxycholate, and nuclei were discarded after a 7-min 15,000g centrifugation. The cytoplasmic optical density (OD) at 260 nm was assessed, 300µg of RNA were resuspended in a final volume of 500 μl and loaded on a 5 to 50% sucrose gradient [20 mM Hepes (pH 7.6), 100 mM KCl, 5 mM MgCl2, 10μg/mlM cycloheximide, 1/10 protease inhibitors, and RNase inhibitor (10 U/ml)] and ultracentrifuged for 2 hours at 36,000 rpm at 4°C in a Beckman SW41Ti rotor. Absorbance at 254nm of the content of the ultracentrifuge tube was measured from top to bottom using a UA-6 UV/VIS detector and 500µL fractions were collected automatically by the machine.

**Web platforms and software**

**STRING** https://string-db.org/cgi/ (44)

**BioGRID** https://thebiogrid.org, Version 4.4.244 (30)

**VENN** diagrams were realized on the with the platform Bioinformatics & Evolutionary Genomics https://bioinformatics.psb.ugent.be/webtools/Venn/

**AIUPred** was used to identify Intrinsically Disordered Protein Regions (65)

**KEGG** https://www.genome.jp/kegg/ (41)

**GO** https://geneontology.org/ (42)

**ClusterProfiler R package** (43)

## Data and materials availability

All data are available in the main text or in the supplementary materials.

## Author contributions

Conceptualization: VG, FR, FMC, AC, JiBa

Investigation: VG, FMC, BM, EFG, SU, AHL, JiBa

Data analysis: VG, FR, JB, MLG, SU, JiBa

Writing: VG and FR

Review & Editing: All authors contributed

---

**Supplementary Figure 1.**
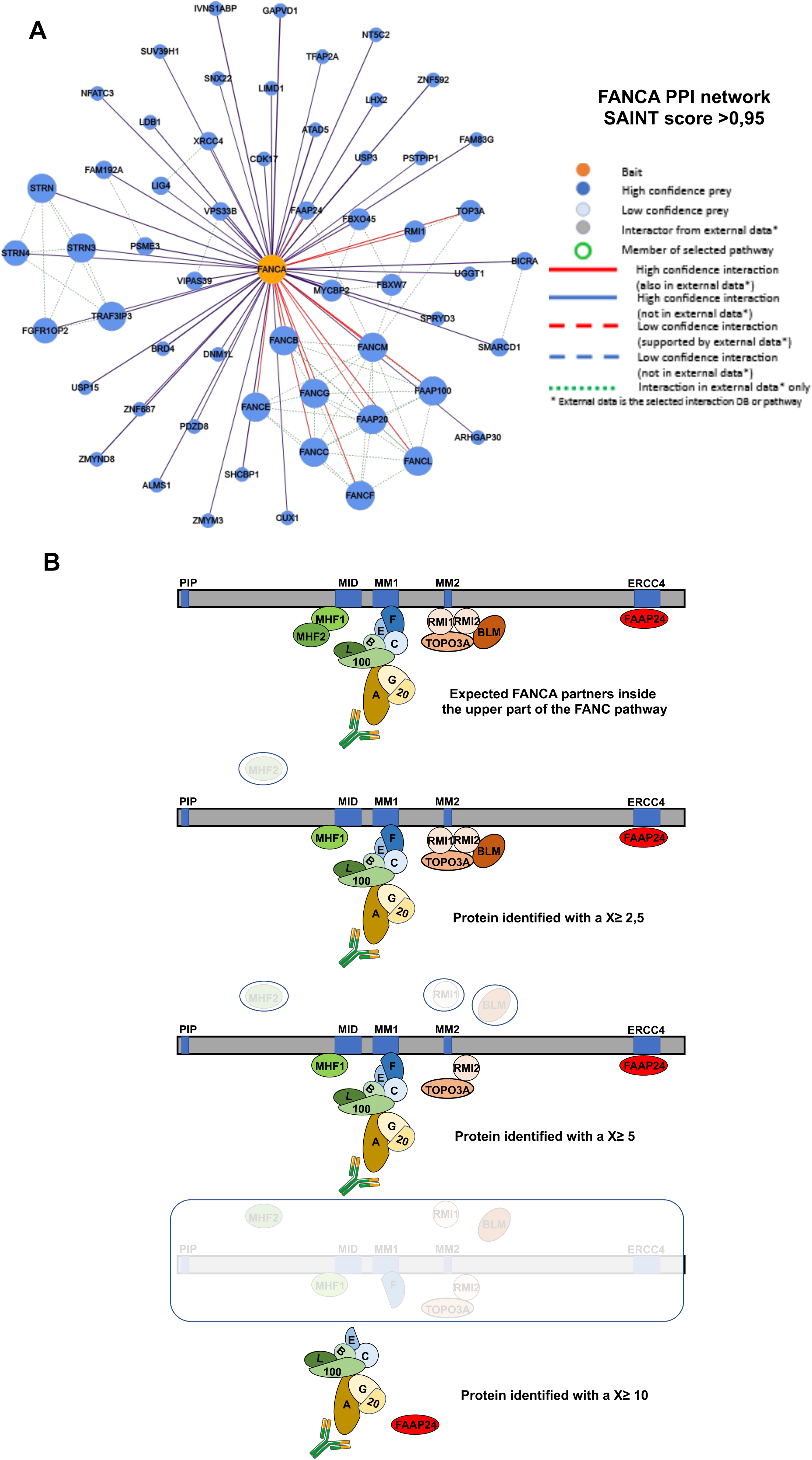
(A) FANCA protein-protein network as determined by SAINT analysis of proteins (SAINT score >0.95) co-immunoprecipitated with FANCA in HSC93 cells. **(B)** Progressive loss of FANCA and FANCM preys in the upper part of the FANC/BRC pathways as function of our selectively levels.

**Supplementary Figure 2.**
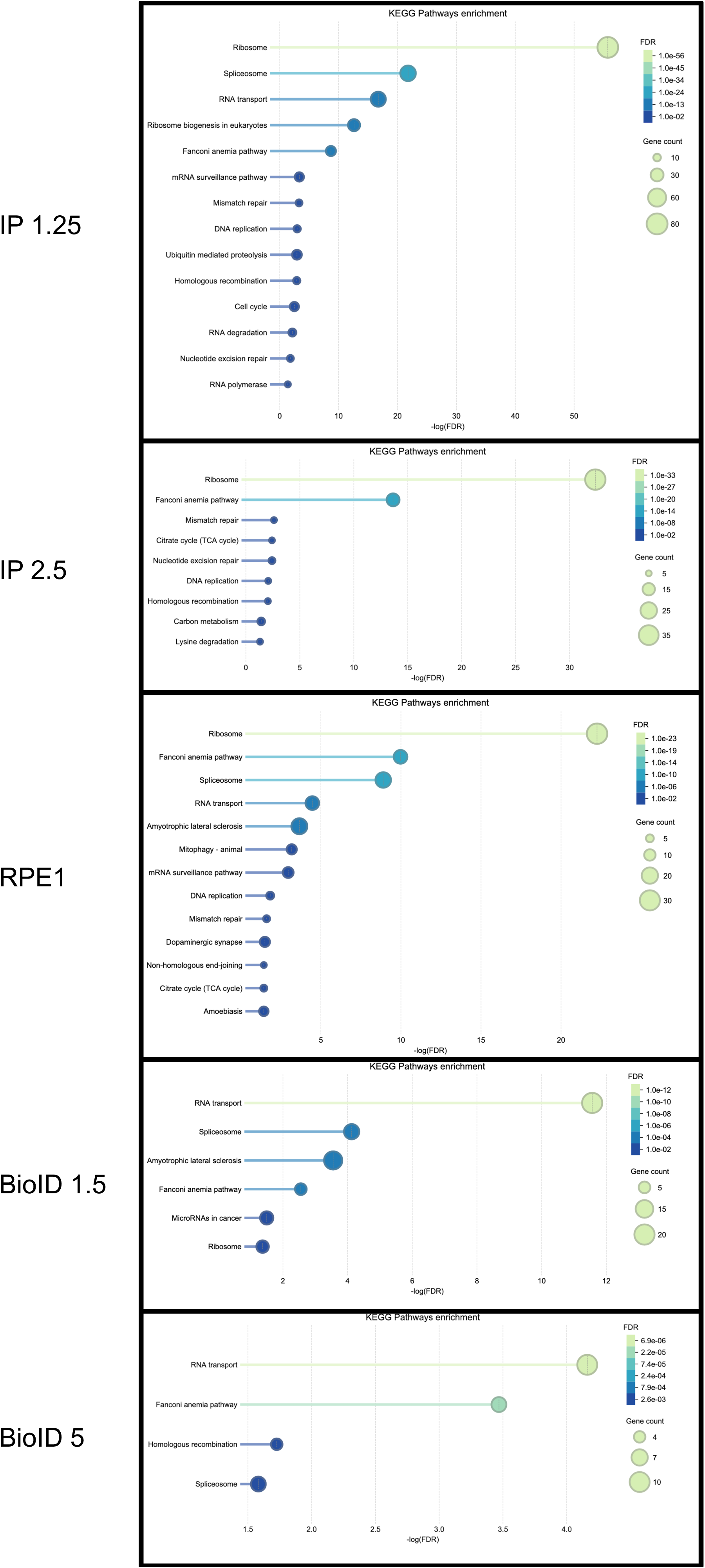
KEGG enriched functional terms as identified by STRING in **X**=1.25 and **X**=2.5, RPE1, BioID>1.5 and BioID>5.

**Supplementary Figure 3.**
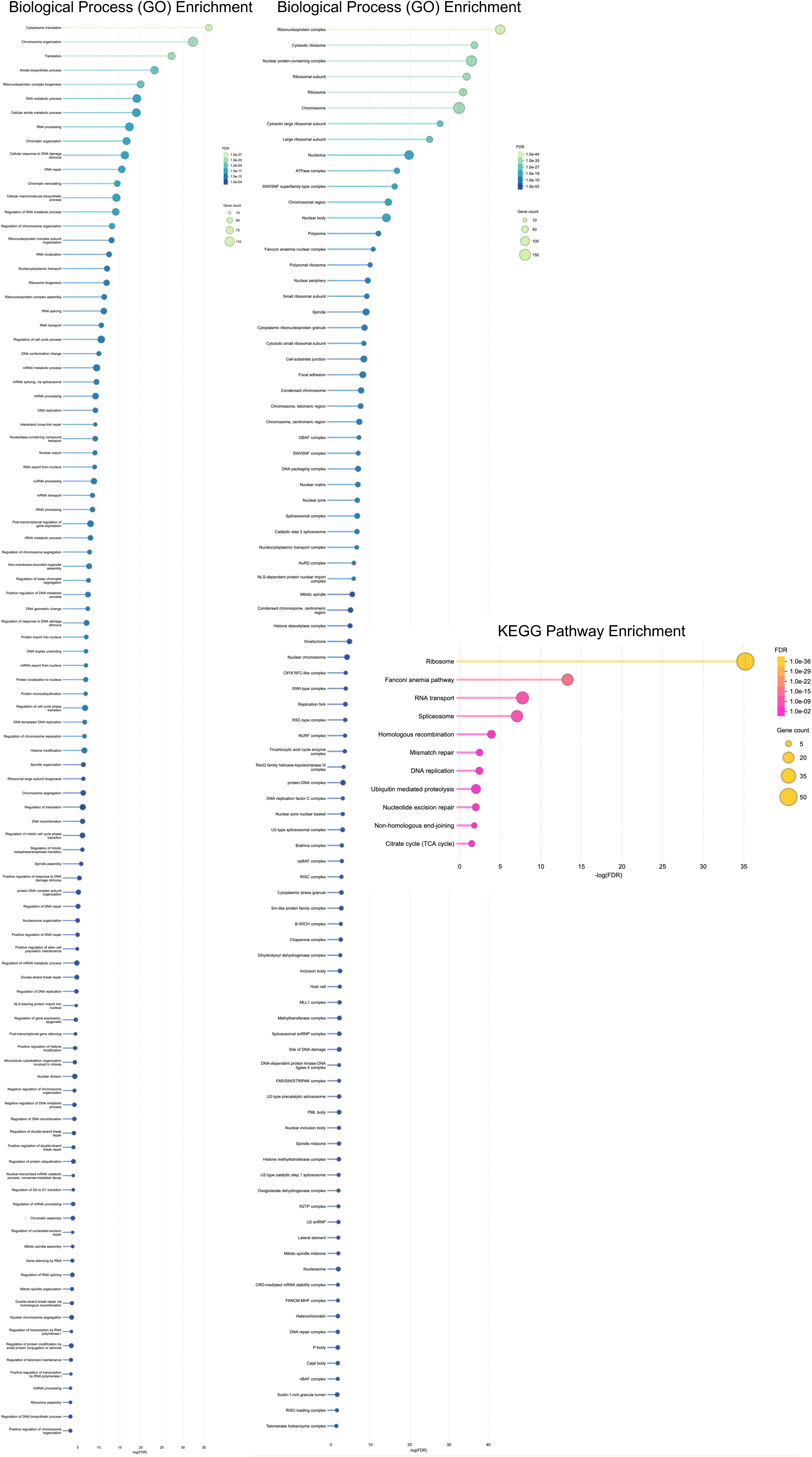
KEGG, GO_BP and GO_CC enriched functional terms as identified by STRING in the AC_2 lists of FANCA potential partners.

**Supplementary Figure 4.**
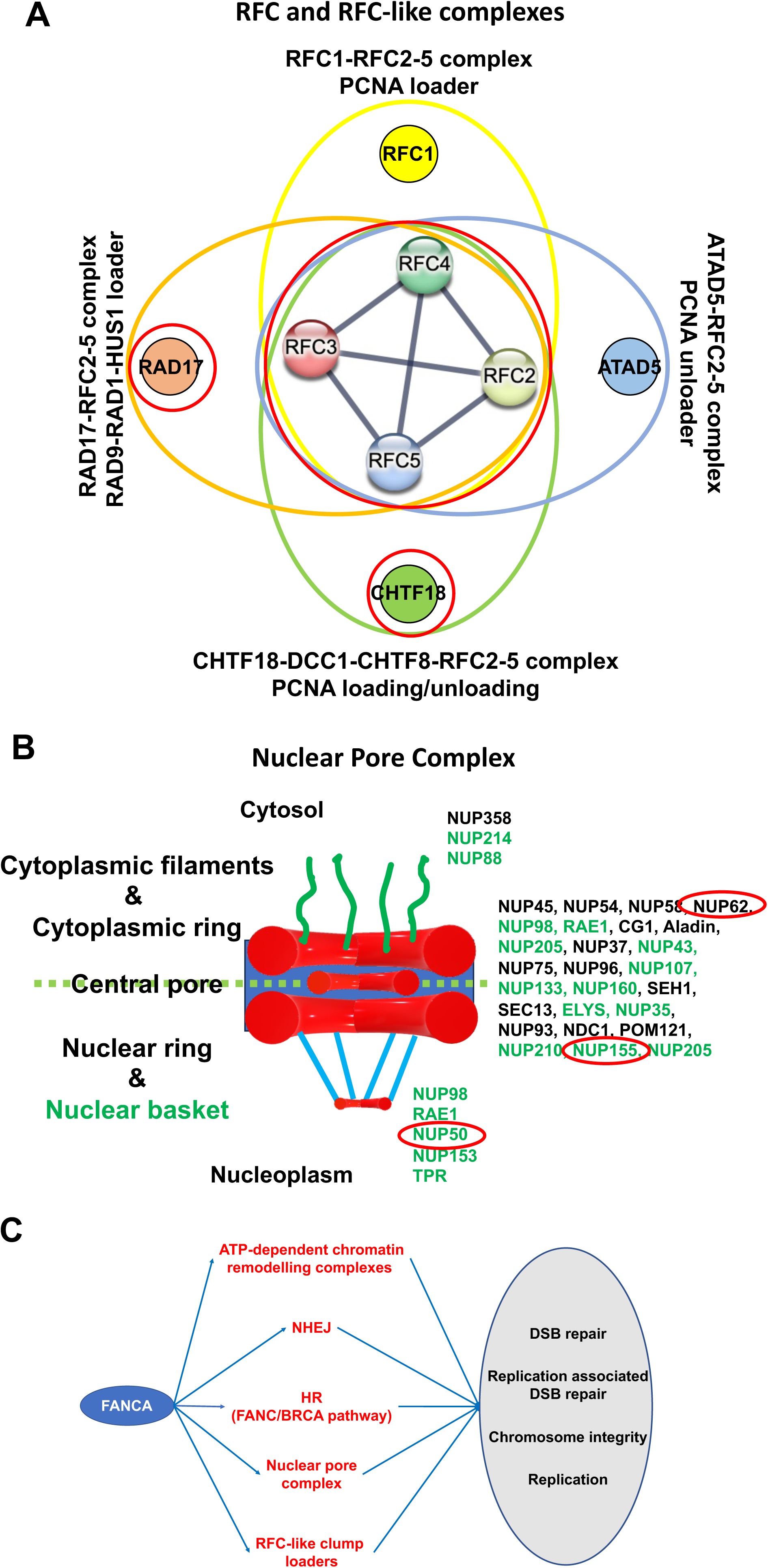
FANCA co-immunoprecipitated or biotinylated preys associated to RFC and RFC-like complexes. The red circle indicates proteins previously identified in the literature. **(A)** FANCA preys associated to the Nuclear Pore Complex reported in green. The red circle indicates proteins previously identified in the literature. **(B)** FANCA links in genome integrity maintenance.

**Supplementary Figure 5.**
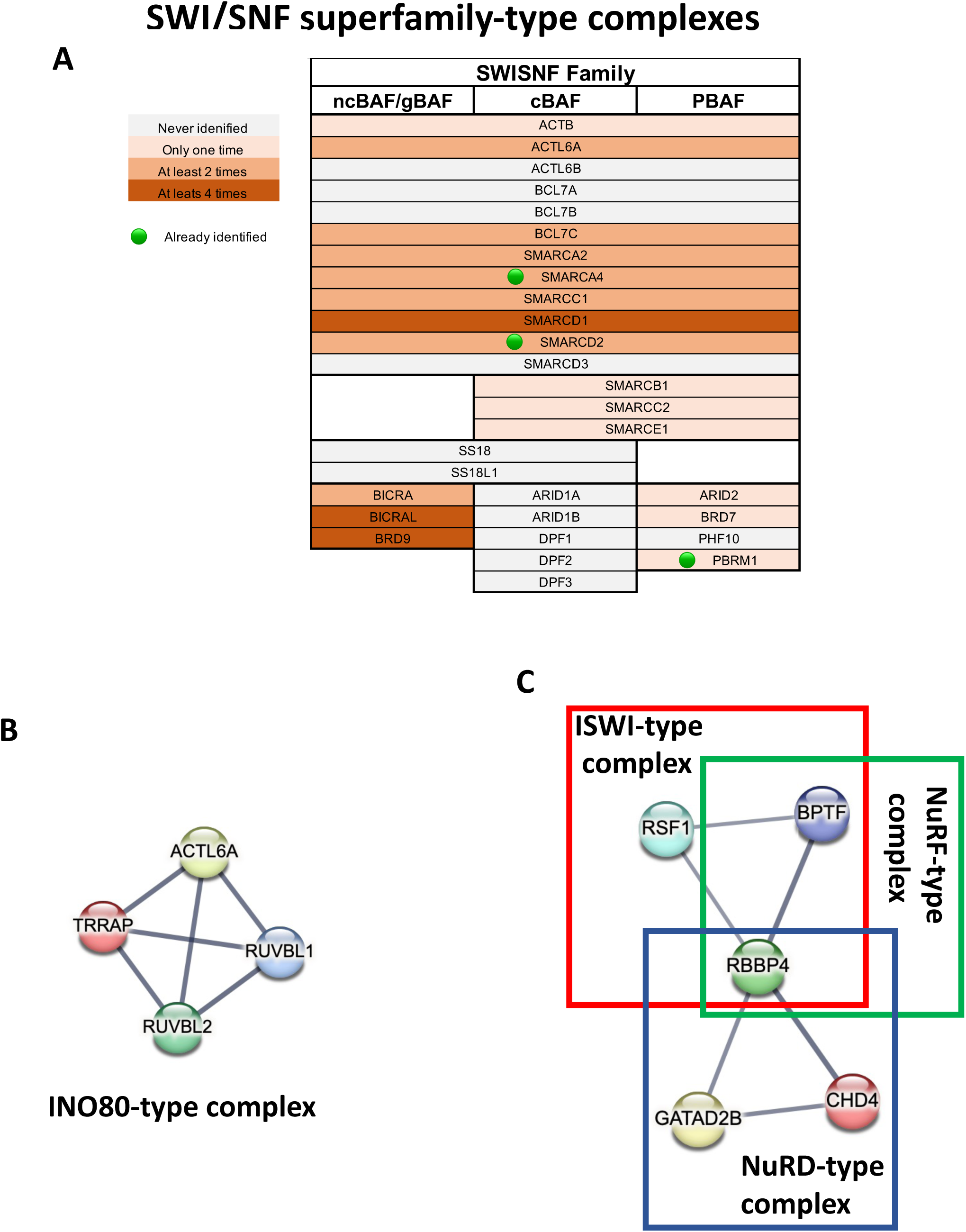
Components of SWI/SNF superfamily complexes identified as FANCA prey. The small green bullet indicates proteins already identified in literature. **(A)** STRING proposed a clustering of FANCA preys associated with the INO80-like complex. **(B)** STRING proposed a clustering of FANCA preys associated with the ISWI-, NuRF-, and NuRD-like complexes.

**Supplementary Figure 6.**
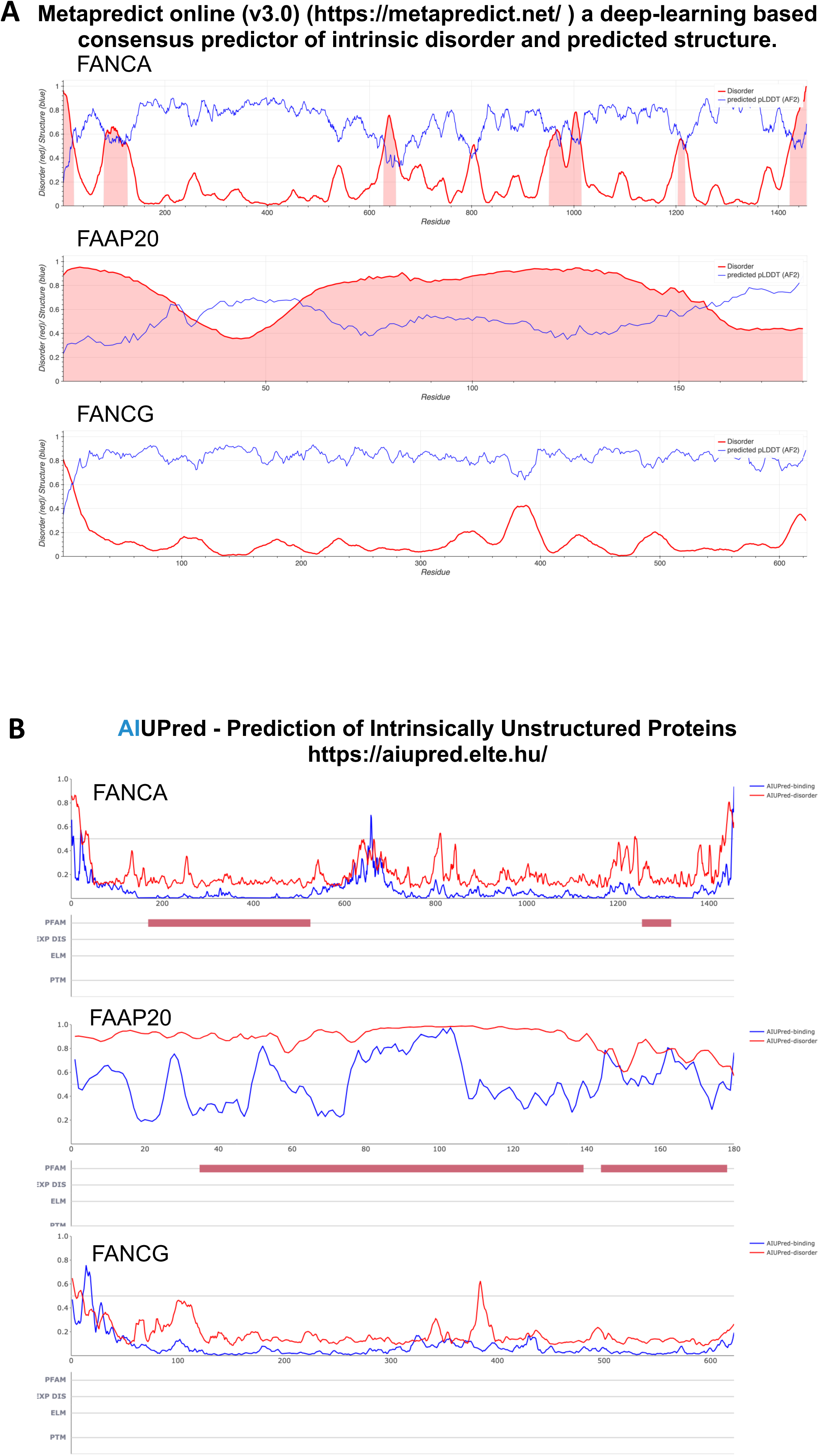
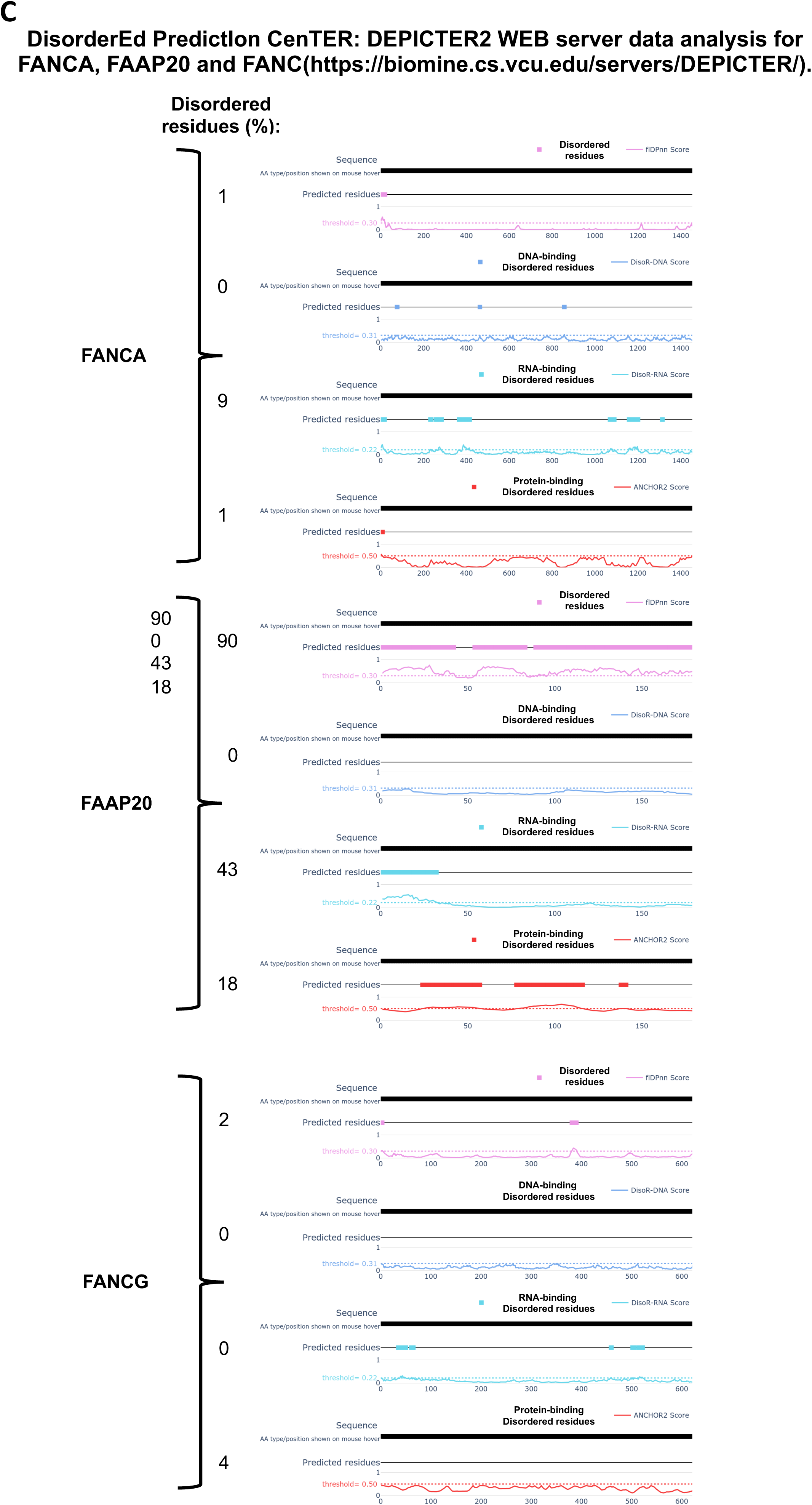
(A-C) Intrinsically Disordered Regions (IDRs) and Structured Regions in FANCA, FAAP20 and FANCG as predicted by Metapredict (v3.0) **(A)**, AIUPRED **(B)** and DEPICTER2 **(C)** WEB sites.

## SUPPLEMENTAL TABLES

**Supplemental Table ST1.**

**Sheet 1.** Mass Spectrometry data from the four independent FANCA immunoprecipitations in HSC93 FANCA-proficient lymphoblasts.

**Sheet 2.** List of FANCA co-immunoprecipitated preys with a SANIT probabilistic score (SP) >0.75.

**Sheet 3.** List of FANCA co-immunoprecipitated preys with a SANIT probabilistic score (SP) >0.95.

**Sheet 4 - 6.** KEGG, GO_Biological Process (BP) and GO_Cellular Components (CC) enrichment analysis of the list of FANCA co-immunoprecipitated preys with a SANIT probabilistic score (SP) >0.75.

**Sheet 7 - 9.** KEGG, GO_Biological Process (BP) and GO_Cellular Components (CC) enrichment analysis of the list of FANCA co-immunoprecipitated preys with a SANIT probabilistic score (SP) >0.95.

**Supplemental Table ST2.**

**Sheet 1.** List of FANCA co-immunoprecipitated preys with a **X**=1.25. **Sheet 2.** List of FANCA co-immunoprecipitated preys with a **X**=2.5. **Sheet 3.** List of FANCA co-immunoprecipitated preys with a **X**=5.

**Sheet 4.** List of FANCA co-immunoprecipitated preys with a **X**=10.

**Sheet 5 - 8.** KEGG enrichment analysis of the list of FANCA co-immunoprecipitated preys with a **X**=1.25, 2.5, 5 and 10, respectively.

**Sheet 9 - 12.** GO_Biological Process (BP) enrichment analysis of the list of FANCA co-immunoprecipitated preys with a **X**=1.25, 2.5, 5 and 10, respectively.

**Sheet 13 - 16.** GO_Cellular Components (CC) enrichment analysis of the list of FANCA co-immunoprecipitated preys with a **X**=1.25, 2.5, 5 and 10, respectively.

**Supplemental Table ST3.**

**Sheet 1.** Datasets of FANCA partners at different selectively level obtained following the immunoprecipitation of FANCA in: HSC93 calculated by the SAINT algorithm or by our in-house filter (green); and, RPE1 cells (orange); and by the BioID approach (blue).

**Sheet 2.** Proteins identified as potential FANCA partners in Lagundžin et al., PlosOne, 2019 following FANCA IP and a SAINT score SP>0,9 (orange); reported in the BioGRID database (Release 4.4.240); identified following different approaches in Reuter et al., Exp Cell Res, 2003; Gueiderikh et al., Sci Adv., 2021; Qian et al., PlosOne, 2013 (light blue); full list of 316 FANCA preys identified in literature (brown); common proteins in Lagundžin et al. (Plo sOne, 2019) & BioGRID (white). KEGG enriched functional terms identified by STRING based analysis in Lagundžin et al., Plo sOne, 2019 (Top) and BioGRID database (Bottom).

**Sheet 3.** Common proteins associated to FANCA between the literature list of 316 preys and our lists at different selectivity levels.

**Sheet 4.** Lists of common FANCA partners obtained comparing our several datasets.

**Sheet 5.** Final lists (additional catalogues, ACs) of FANCA preys calculated on the basis of their presence in more of the previous described datasets used for the building of our proposed multilayered FANCA PPI network in Figure 6).

**Sheet 6 - 12.** FANCA-associated KEGG and GO_CC sub-networks: the FANC pathway; the NHEJ pathway; the ribosomal/translational network; the spliceosome/RNA splicing network; the replication network; the SWI/SNF network; and, the Nuclear Pore Complex/nucleocytoplasmic transport network, respectively for Sheet 6 to 12. For each subnetwork the ID of each protein and their presence in the previous AC are reported

**Supplemental Table ST4.**

**Sheet 1.** Mass Spectrometry data from FANCA immunoprecipitation in RPE1 cells.

**Sheet 2 - 4** KEGG, GO_Biological Process (BP) and GO_Cellular Components (CC) enrichment analysis of the list of FANCA co-immunoprecipitated preys in RPE1 as obtained by STRING analysis.

**Supplemental Table ST5.**

**Sheet 1.** Mass Spectrometry data from FANCA-BirA-mediated biotinylation the FANCA proxilome.

**Sheet 2 - 3** KEGG, GO_Biological Process (BP) and GO_Cellular Components (CC) enrichment analysis of the list of FANCA biotinylated targets at IR>1.5 and IR>5, respectively.

**Supplemental Table ST6.**

KEGG enriched terms, as identified by STRING, in IP **X**=1.25, **X**=2.5, RPE1, BioID>1.5 and BioID>5.

**Supplemental Table ST7.**

Quantification of the WB for Figure 8.

**Sheet 1.** Polysome fractions in Exp1 were analyzed twice by WB (data from one WB are presented in Figure 8B and 8C). The mean of the two measures was used for Figure 8D. **Sheet 2.** Polysome fractions in Exp2 were analyzed once by WB (not shown).

**Sheet 3 and 4.** Data from previous sheets to draw Figure 8E and 8F.

